# Engineered translation release factor 1 suppresses disease-causing nonsense mutations and enables multi-site nonstandard amino acids incorporation

**DOI:** 10.64898/2025.12.24.695955

**Authors:** Yuting Chen, Meilin Tian, Zhu Zhu, Ya Qian, Yichong Xu, Peng Gong, George Church, Chenli Liu

## Abstract

Nonsense mutations convert sense codons into premature termination codons (PTC), resulting in early termination of translation of mRNAs, underlie ∼11% of human genetic diseases. Restoring translation of genes carrying nonsense mutations remains a major therapeutic challenge. Here we present OPENER (Overexpression of Protein-engineered eRF1 for Nonsense Elision and Readthrough), a strategy to efficiently suppress nonsense mutations based on engineered variants of the eukaryotic translation termination factor eRF1. Through saturation mutagenesis of the human eRF1 N-domain, we identified variants, notably eRF1^S77E^, that promote efficient readthrough of all three PTCs (UAG, UGA, UAA) and rescue 12 pathogenic PTCs reporters in multiple disease contexts. Systemic delivery of OPENER-S77E via AAV8 in a Duchenne muscular dystrophy mouse model (*Dmd*^Q995X^) restored dystrophin expression, improved muscle function, and ameliorated pathology without detected toxicity. Furthermore, when combined with orthogonal engineered aminoacyl-tRNA synthetase/tRNA pairs, OPENER-S77E enabled efficient, multi-site incorporation of nonstandard amino acids (BocK and AzF) into a single protein in mammalian cells, achieving efficiency of 27.77% and 19.83%, respectively. The incorporation efficiency of BocK was further enhanced to 32.80% in conditional eRF1-knockout cells. OPENER thus provides a platform for treating nonsense-mediated diseases and expands the synthetic biology toolkit for precise protein engineering.

## Introduction

Nonsense mutations introduce premature termination codons (PTCs) in mRNAs, leading to truncated and nonfunctional proteins. These mutations account for 11% of pathogenic mutations, e.g. Cystic fibrosis, Hurler syndrome and Duchenne muscular dystrophy(*1, 2*). Several therapeutic strategies have been developed to treat PTC-related rare genetic diseases(*3*). For example, gene replacement(*4*), gene editing(*5*), base editing(*6, 7*), RNA editing(*8, 9*) and pseudouridylation(*10, 11*) have been used to correct PTCs, but each faces limitations including low efficiency, high off-target, strong immunogenicity and limited delivery capacity for large gene size. Moreover, the translational readthrough of PTCs has shown promise in restoring full-length protein expression. Aminoglycosides (G418, paromomycin and gentamicin) and non-aminoglycosides (PTC124) show some efficiency for PTC suppression in vitro and animal models (*12–14*), but their clinical benefit remains limited due to low efficiency, global toxicity, non-specific insertion of random amino acids, and off-target readthrough at natural stop codons. Engineered suppressor tRNAs have been shown to suppress nonsense mutants in vivo(*15–17*), but often require overexpression to potentially toxic levels to achieve therapeutic efficacy. Prime editing-installed suppressor tRNAs can rescue nonsense mutations in a disease-agnostic manner(*18*), which may require unnecessary long-term exposure to editing agents, leading off-target hits and immunogenicity. To tackle such a challenge, we are focusing on reprogramming translation termination factor, human eukaryotic release factor 1 (eRF1) encoded by *ETF1* gene(*19*). Human eRF1 as a single omnipotent factor is responsible for recognition of all three stop codons (UAG, UGA, and UAA) and peptidyl-tRNA hydrolysis, acting in the eRF1-eRF3-GTP ternary complex in translation termination when a stop codon on the mRNA enters the A site of ribosomes(*20–23*). Based on mutagenesis and structure analyses, human eRF1 has a tRNA-shaped protein factor composed of three domains(*24*), and the N-domain of human eRF1 is responsible for stop codon recognition(*25, 26*). This domain contains the highly conserved NIKS motif (residues 61-64)(*27*), YxCxxxF motif (residues 125-131) and the GTS loop (residues 31-33)(*28, 29*), all of which have been implicated in stop codon recognition. However, previous eRF1 mutagenesis studies have focused on limited amino acid sites(*30–32*) and systematic understanding of human eRF1 recognition domain remains incomplete. While engineered tRNA synthase-tRNA pairs have been reported to genetically encode nonstandard amino acids (nsAAs) in many species(*33–35*), their efficiency has been limited, so far, in mammalian cells and animals(*36*), even with overexpression of eRF1^E55D^ and other eRF1 variants, only modest improvements in the incorporation of multiple nsAAs have been reported, with incorporation efficiency of BocK increasing from ∼ 5% to ∼ 12%(*32, 37*).

Although translational readthrough reagents could in principle read through both PTCs and natural termination codons (NTCs), extensive evidence indicates that several such agents are generally well tolerated in mammalian cells. This tolerance arises from multiple biological mechanisms that have been well documented in previous studies (*12, 18*). Meanwhile small-molecule-mediated eRF1 depletion enhances readthrough of PTCs in human cells(*38, 39*). We therefore hypothesized that targeted weakening of eRF1’s termination fidelity, without ablating its essential function at NTCs, could provide a general strategy for promoting PTC readthrough. Here, we describe a comprehensive saturation mutagenesis screen of the eRF1 N-domain, identifying gain-of-function variants that promote efficient PTC suppression. We term this approach OPENER (Overexpression of Protein-engineered eRF1 for Nonsense Elision and Readthrough). We demonstrate its therapeutic potential by rescuing dystrophin expression and function in a mouse model of Duchenne muscular dystrophy (DMD). Moreover, by decoupling OPENER from near-cognate tRNA competition and coupling it to orthogonal translation systems, we repurpose it to significantly enhance the multi-site incorporation of nonstandard amino acids in human cells. This work establishes OPENER as a dual-purpose platform for therapeutic gene function restoration and advanced synthetic biology.

## Results

### Identification of OPENER suppressing nonsense mutants by saturated mutagenesis library screens targeting the N-terminal domain of human eRF1

To determine whether overexpression of engineered human eRF1 can promote premature stop codon readthrough, we performed a saturated mutagenesis screen targeting the N-terminal domain of human eRF1, which is responsibility for stop codon recognition. Moreover, we generated a stable human cell line HEK293T which expressing a sfGFP reporter containing a UAG stop codon (sfGFP^150TAG^) using lentivirus(*32*). In this reporter system, the 150th codon of the sfGFP mRNA was mutated to an amber codon (UAG) and full-length sfGFP green fluorescent protein is expressed in cells when the amber codon is successfully suppressed, allowing quantitative measurement of PTC readthrough by monitoring sfGFP fluorescence. The lentivirus human eRF1 variants library was transduced into HEK293T-sfGFP^150TAG^ reporter cells at low MOI (∼ 0.1) and sfGFP positive single clones were isolated and cultured after flow cytometry (FACS) sorting (Fig. 1A and figs. S1A, S1B). Populations of clones with sfGFP fluorescence were expanded and each clone were collected for next-generation sequencing to identify enriched eRF1 variants (figs. S1C and S1D). For data analysis, we have calculated and ranked human eRF1 variants according to the frequency and proportion of occurrences across different clones. The most recurrent and highly enriched eRF1 variants were selected for further validation (Fig. 1B and table S1).

**Fig. 1.**
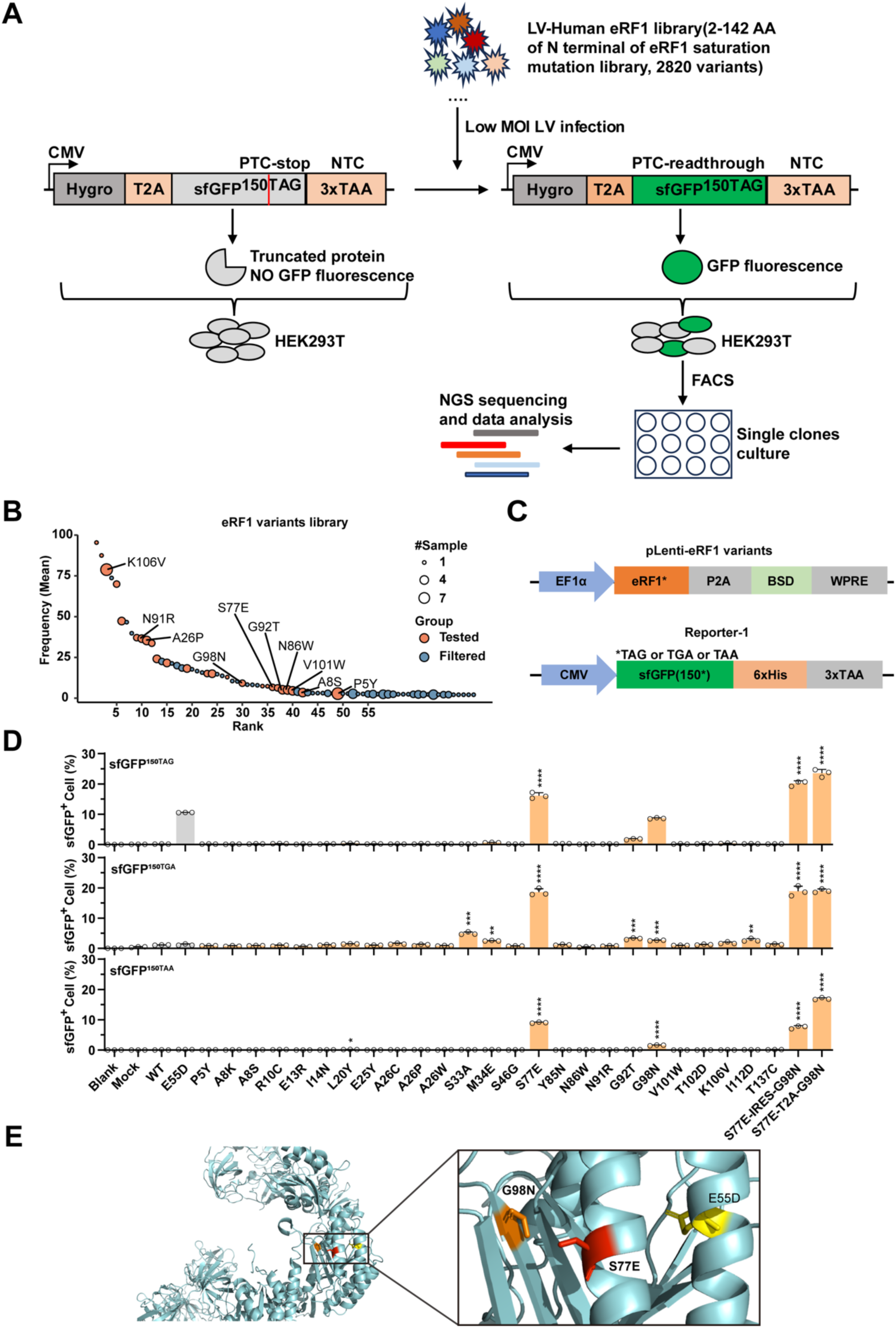
Identification of OPENER suppressing PTCs by a saturated mutagenesis library screen targeting the N-terminal domain of human eRF1. (A) Workflow of the saturated mutagenesis library screening. (B) Frequency distribution of engineered eRF1 variants in single clones. (C) Schematic of the human eRF1 variant constructs and Reporter-1s containing the 150th codon of sfGFP mRNA was mutated to UAG, UGA, UAA, separately. (D) Validation of OPENER in HEK293T cells. Flow analysis was performed 72h after co-transfection of each Reporter-1s constructs with OPENER separately. Bar plot shows the fraction of sfGFP positive cells. Wild-type eRF1 and eRF1^E55D^ served as controls. Data are mean ± SD, n = 3; \**P* < 0.05; \*\**P* < 0.01; \*\*\**P* < 0.001; \*\*\*\**P* < 0.0001, compared to eRF1^E55D^ (two-sided Student′s *t*-test). (E) positions of eRF1^S77E^, eRF1^G98N^ and eRF1^E55D^ mapped to the structure of the N-terminal domain of eRF1 (PDB ID:3E1Y).

To validation of the high screen hits, we co-transfected OPENER-engineered eRF1 variants with sfGFP^150TAG^ into HEK293T separately (Fig. 1C). Although eRF1^E55D^ was not identified in our screens and has not been reported to directly promote PTC readthrough. eRF1^E55D^ was included in the validation experiments because overexpression of eRF1^E55D^ is known to promote nsAAs incorporation(*32*), Flow cytometry analysis showed that the fractions of sfGFP positive cells were 16.13% (OPENER-S77E), 8.66% (OPENER-G98N), compared to 10.57% (eRF1^E55D^). Notably, combining OPENER-S77E with OPENER-G98N further enhanced readthrough efficiency to 23.57% (OPENER-S77E-T2A-G98N) and 20.20% (OPENER-S77E-IRES-G98N). These data demonstrate that OPENER-S77E alone and combined with OPENER-G98N efficiently suppresses the UAG stop codon (Figs. 1D and 1E).

To test whether OPENER can induce readthrough of UGA and UAA codons, we constructed another two reporters, sfGFP^150TGA^ and sfGFP^150TAA^, and co-transfected OPENER with each reporter into HEK293T separately. For the UGA readthrough, the fractions of sfGFP positive cells were 18.57% (OPENER-S77E), 19.03% (OPENER-S77E-T2A-G98N), and 18.90% (OPENER-S77E-IRES-G98N), and 4.97% (OPENER-S33A), 2.51% (OPENER-M34E), 3.20% (OPENER-G92T), 2.63% (OPENER-G98N), 2.77% (OPENER-I112D), compared to 1.22% (eRF1^E55D^) (Fig. 1D). For the UAA codon readthrough, the fractions of sfGFP positive cells were 9.06% (OPENER-S77E), 1.58% (OPENER-G98N), 16.93% (OPENER-S77E-T2A-G98N), and 7.57% (OPENER-S77E-IRES-G98N), compared to 0.14% (eRF1^E55D^) (Fig. 1D). The geometric mean fluorescence intensity of sfGFP was also quantified and showed a trend consistent with the significant increase in the fractions of sfGFP positive cells (figs. S2A-C). Altogether, we conclude that OPENER-S77E and OPENER-S77E-T2A-G98N can enhance readthrough all three stop codons. OPENER-S77E-T2A-G98N showed the highest readthrough at the amber codon (UAG), followed by the opal codon (UGA) and ochre codon (UAA), whereas OPENER-S77E showed the highest readthrough at the UGA, followed by UAG and UAA.

### OPENER rescue protein expression of PTCs reporters

We further measured PTC readthrough events of OPENER using a high-content imaging system and western blotting analysis. sfGFP fluorescence and full-length sfGFP protein were observed when co-transfection of OPENER with Reporter-1s containing UAG, UGA, or UAA codons into HEK293T cells, respectively. These results confirmed robust expression of full-length sfGFP and the readthrough efficiency were consistent with flow cytometry analysis (Figs. 2A-D). To assess the general applicability of OPENER across cell types, we used an additional murine cell line, Neuro-2a. In this cell line, we observed that OPENER significantly enhances PTC readthrough events using Reporter-1s, and the trend in readthrough efficiency are consistent with that observed in HEK293T cells, although efficiencies were modestly lower, likely due to differences in transfection efficiency (figs. S3A-C). Together, these data suggest that OPENER is a broadly applicable tool for suppressing PTCs in different mammalian cells.

**Fig. 2.**
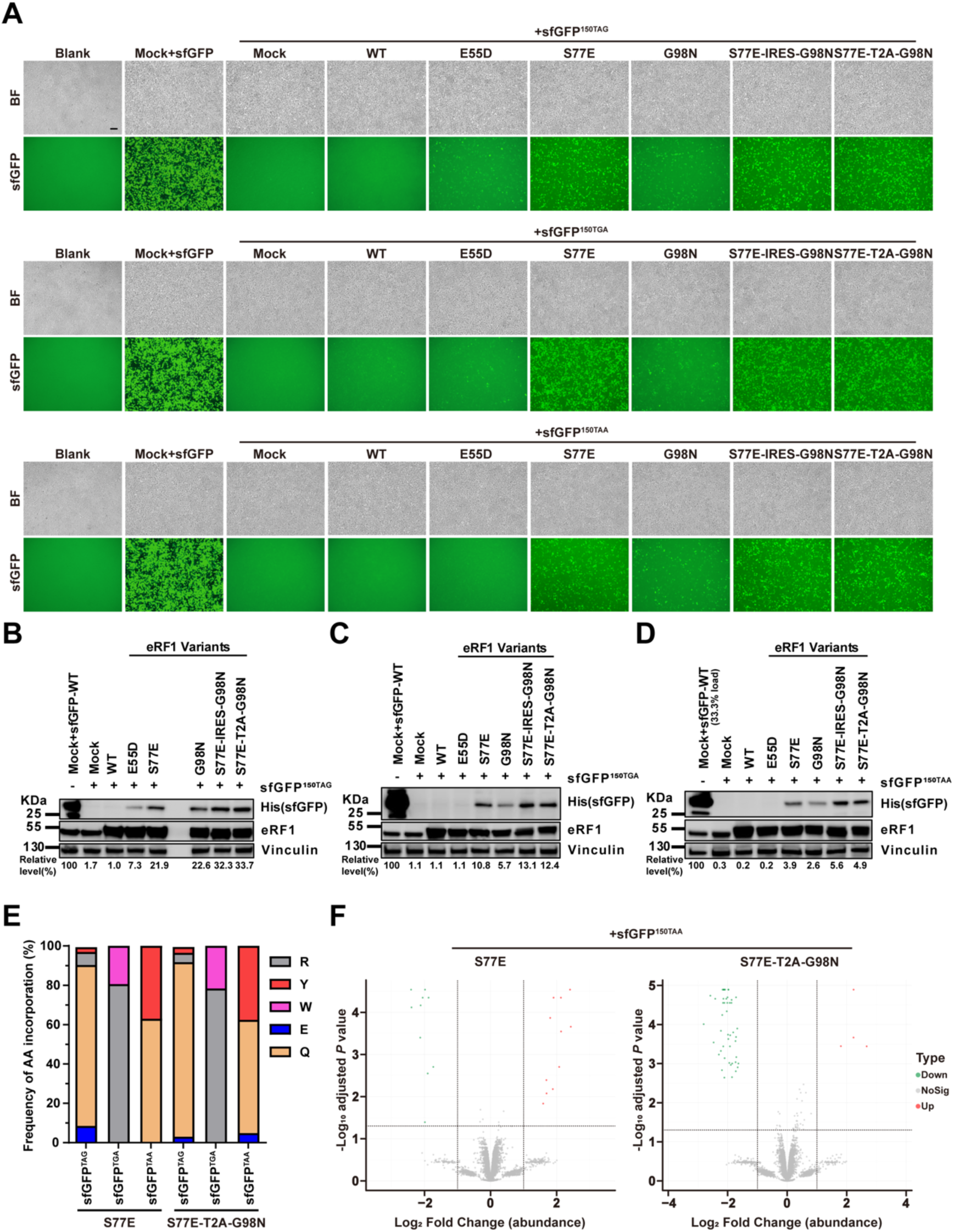
OPENER can rescue protein expression of PTCs reporters. (A) Representative fluorescence images of cells showing the expression levels of sfGFP. Scale bars, 100 μm. (B-D) Western blotting analysis showing the expression levels of full-length sfGFP proteins when co-transfection of OPENER with sfGFP^150TAG^ (B), sfGFP^150TGA^ (C) and sfGFP^150TAA^ reporter (D). For semi-quantification, the integrated optical density (IOD) of sfGFP bands was normalized to that of sfGFP-WT after load correction. Normalized IOD values are shown underneath the blot. WT, wild-type. (E) Amino acid incorporated at the 150th residue of sfGFP were determined by mass spectrometry analysis of HA-tagged sfGFP harvested 72h after co-transfection of three Reporter-1s with OPENER-S77E and OPENER-S77E-T2A-G98N separately. (F) Protein-level analysis of whole proteome mass spectrometry data from sfGFP positive cells sorted by FACS after co-transfection of OPENER-S77E and OPENER-S77E-T2A-G98N with sfGFP^TAA^ reporter separately. Log_2_ fold change in protein abundance compared to sfGFP-WT control cells. Dashed lines indicate an adjusted *p*-value of 0.05 on the y-axis and log_2_ fold change in abundance of ±1 compared with control cells.

To identify which amino acids are incorporated during the OPENER mediate readthrough of PTC sites in Reporter-1s in human cells, we carried out immunoprecipitations to isolate the full-length sfGFP proteins and subjected the proteins to mass spectrometry after co-transfection of OPENER with Reporter-1s into HEK293T. LC-MS/MS analysis showed that the amino acids incorporated by OPENER-S77E-mediated stop codons suppression were (1) Gln (62.96%) and Tyr (37.04%) at UAA; (2) predominantly Gln (81.79%), with lower frequency of Tyr, Arg, and Glu at UAG; and (3) Arg (80.57%) and Trp (19.43%) at UGA. OPENER-S77E-T2A-G98N-mediated stop codons suppression were (1) predominantly Gln (57.57%) and Tyr (37.53%), with lower frequency of Glu at UAA; (2) predominantly Gln (88.76%), with a lower frequency of Tyr, Arg, and Glu at UAG; and (3) Arg (78.38%) and Trp (21.62%) at UGA (Fig. 2E and table S2). This incorporation pattern is very similar to amino acids incorporation preference reported for basal or drug-induced readthrough(*40*), as well as RNA pseudouridylation-dependent readthrough(*10*).

### Effect of OPENER on the proteome

The aminoglycosides such as G418(*41*) and some sup-tRNAs(*42*) was widely reported that can induce global toxicity for cells when addition of them to cells, which poses a safety concern in therapeutic settings. To evaluate potential effect of OPENER for proteome, we performed label-free quantitative mass spectrometry on trypsin-digested total protein lysates from both positive control (wild-type sfGFP) and sfGFP positive cells were collected by FACS sorting after co-transfection of OPENER mediate readthrough of PTC sites in Reporter-1s separately. Mass spectrometry data showing the only few proteins that were significantly (adjusted *P* value < 0.05, more than 2-fold change) different in abundance between control samples and OPENER samples (Fig. 2F and figs. S4A, S4B).

### Quantifying readthrough of human pathogenic PTCs by OPENER

To test whether OPENER can readthrough of diverse human pathogenic PTCs and quantify OPENER induced readthrough, we constructed additional Reporter-2s. Because the local sequence contexts of PTCs has been reported to influence the readthrough efficiency of PTC by drugs and other factors(*41*), and each pathogenic PTC was cloned together with 60 nucleotides of surrounding sequence into a dual fluorescent protein reporter, in which an upstream mCherry protein serves as normalization controls and readthrough causes expression of a downstream green fluorescent protein EGFP (Fig. 3A).We selected 12 clinically relevant PTCs containing three different stop codons in *IDUA, CFTR, DMD* and *RPE65* genes(*43*), associated with four common genetic diseases, Hurler syndrome, Cystic fibrosis, Duchenne muscular dystrophy, and Leber congenital amaurosis, respectively. EGFP signal was evaluated by a high-content imaging system and quantified by flow cytometry and western blotting comparison with the positive control group. We found EGFP fluorescence across all PTCs after co-transfection of OPENER-S77E and OPENER-S77E-T2A-G98N and Reporter-2s separately in HEK293 cells (Fig. 3B). Flow cytometry analysis showed that OPENER-S77E and OPENER-S77E-T2A-G98N can efficiently readthrough of all 12 PTCs, and the relative fractions of EGFP positive cells were from 28.94% to 68.00% (OPENER-S77E), and from 41.72% to 80.23% (OPENER-S77E-T2A-G98N) (Fig. 3C and figs. S5A, S5B). Western blotting showed that the OPENER-S77E and OPENER-S77E-T2A-G98N restored all full-length Reporter-2s protein expression (Fig. 3D), and efficiency of OPENER were consistent with those observed by flow cytometry analysis. Together, we found that the OPENER-S77E and OPENER-S77E-T2A-G98N efficiently suppress all 12 pathogenic PTCs across multiple disease-associated sequence contexts in reporters, highlighting their broad applicability for readthrough of diverse nonsense mutations in human genes.

**Fig. 3.**
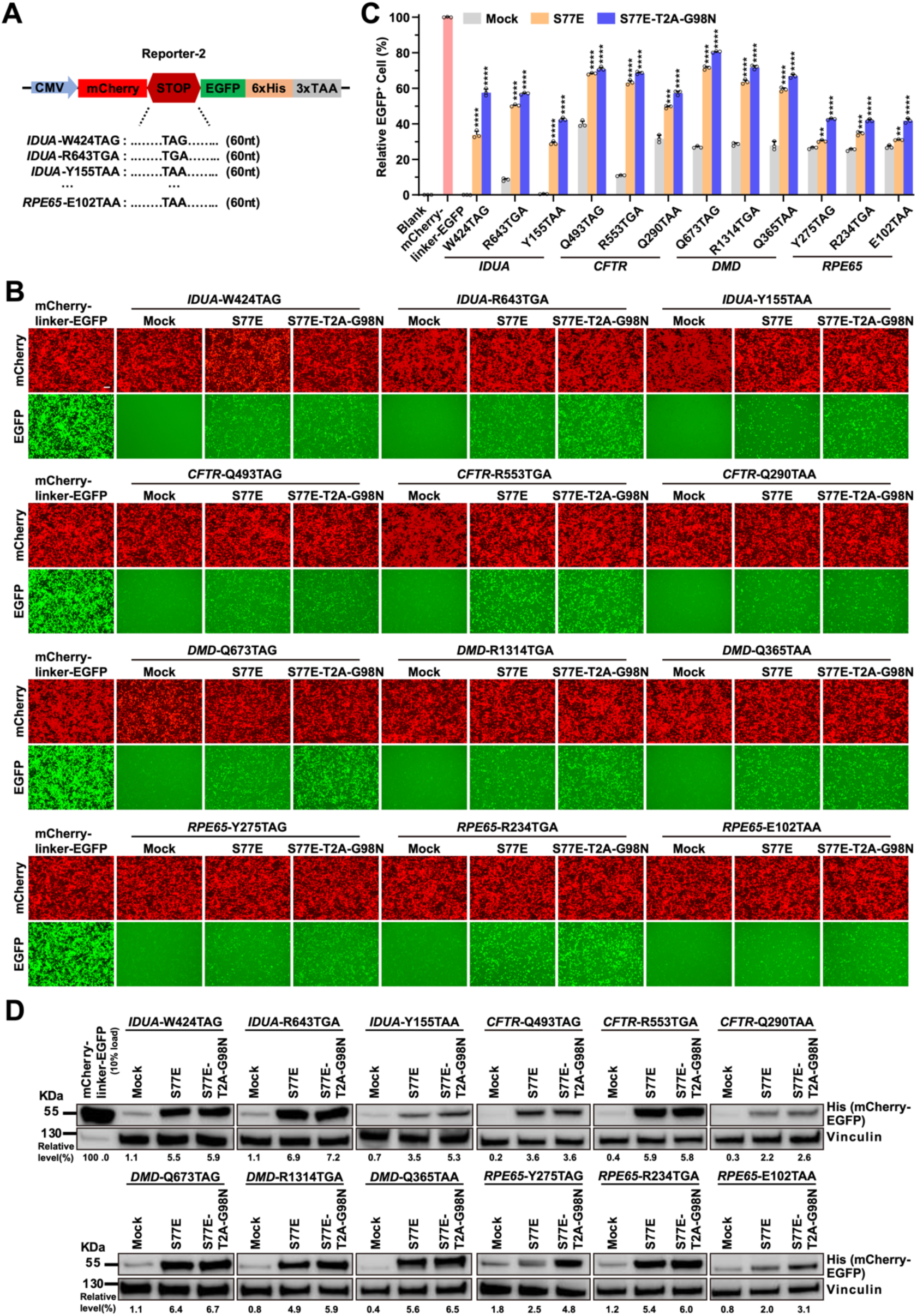
OPENER suppress human pathogenic PTCs in reporters. (A) Schematic of the Reporter-2s constructs. Each pathogenic PTC with 60 nucleotides (nts) of surrounding sequence context was inserted into a dual fluorescent protein reporter. Full sequences of 60 nucleotides are shown in Supplementary fig.5A. (B) Representative fluorescence images of cells showing the expression levels of mCherry-EGFP fusion proteins, 72h after OPENER-S77E and OPENER-S77E-T2A-G98N with Reporter-2s constructs were transfected into HEK293T, respectively. Scale bars, 100 μm. (C) Flow cytometry analysis performed, 72h after co-transfection of OPENER-S77E and OPENER-S77E-T2A-G98N and Reporter-2s constructs were transfected into HEK293T, respectively. Bar plot shows the relative fraction of EGFP positive cells normalized to that of the mCherry-linker-EGFP. Data are represented as the mean ± SD, n = 3; \**P* < 0.05; \*\**P* < 0.01; \*\*\**P* < 0.001; \*\*\*\**P* < 0.0001, compared to Mock (two-sided Student′s *t*-test). (D) Western blotting analysis showing the expression levels of full-length mCherry-EGFP fusion proteins generated by OPENER mediated readthrough of PTCs, 72h after co-transfection of each Reporter-2s with OPENER-S77E and OPENER-S77E-T2A-G98N separately. The integrated optical density (IOD) of mCherry-EGFP fusion proteins bands was normalized to that of the mCherry-linker-EGFP after load correction. Normalized IOD values are shown underneath the blot.

### Restoration of dystrophin in *Dmd*^Q995TAA^ mice by AAV-OPENER

To evaluate the therapeutic potential of AAV-OPENER, we used a mouse model carrying a homozygous nonsense mutation in the *Dmd* gene (*Dmd*^Q995X^, CAA→TAA). This mouse model recapitulates key pathologies features of human DMD, including progressive muscle weakness and a shortened life span(*44–46*). We chose this disease model of *Dmd*^Q995TAA^ mice because nonsense TAA is poorly responsive to PTC-readthrough drugs, and difficult to be editing by DNA and RNA base editors. We first cloned 60 nts context sequence of Q995X of *Dmd* gene into Reporter-2, and observed PTC readthrough after co-transfection of AAV-OPENER and Reporter (Fig. 4A and fig. S6A). Flow cytometry analysis showed that efficiency of OPENER-S77E and OPENER-S77E-T2A-G98N were 58.30% and 56.44%, which were higher than that of G418 (25.80%) and Mock (21.90%), and geometric mean fluorescence intensity of EGFP was the highest by OPENER-S77E mediated readthrough (Figs. 4B, 4C and figs. S6B, S6C). We next constructed AAV8 carrying OPENER-S77E, and injected it into *Dmd*^Q995TAA^ mice intravenously to evaluate the in vivo dystrophin restoration (Fig. 4D). 10 weeks after injection, immunostaining showed dystrophin restoration in tibialis anterior muscle, heart and diaphragm by the intravenous AAV-OPENER-S77E treatment (Fig. 4E). Consistent with improved muscle integrity, serum creatine kinase levels in AAV-OPENER-S77E treated *Dmd*^Q995TAA^ mice were substantially decreased at 10 weeks compared with *Dmd*^Q995TAA^ mice (Fig. 4F). Grip strength testing also showed a significant increase in muscle strength of AAV-OPENER treated *Dmd*^Q995TAA^ mice at 10 weeks compared with control mice (Fig. 4G). AAV-OPENER-S77E treatment significantly increased the expression level of *Dmd* mRNA in heart and tibialis anterior muscles (Fig. 4H). Together, these results demonstrate that AAV-OPENER enables efficient in vivo suppression of a disease-causing TAA nonsense mutation and restores dystrophin expression and muscle function in *Dmd*^Q995TAA^ mice, supporting potential and possibility to clinical applications of the AAV-OPENER approach.

**Fig. 4.**
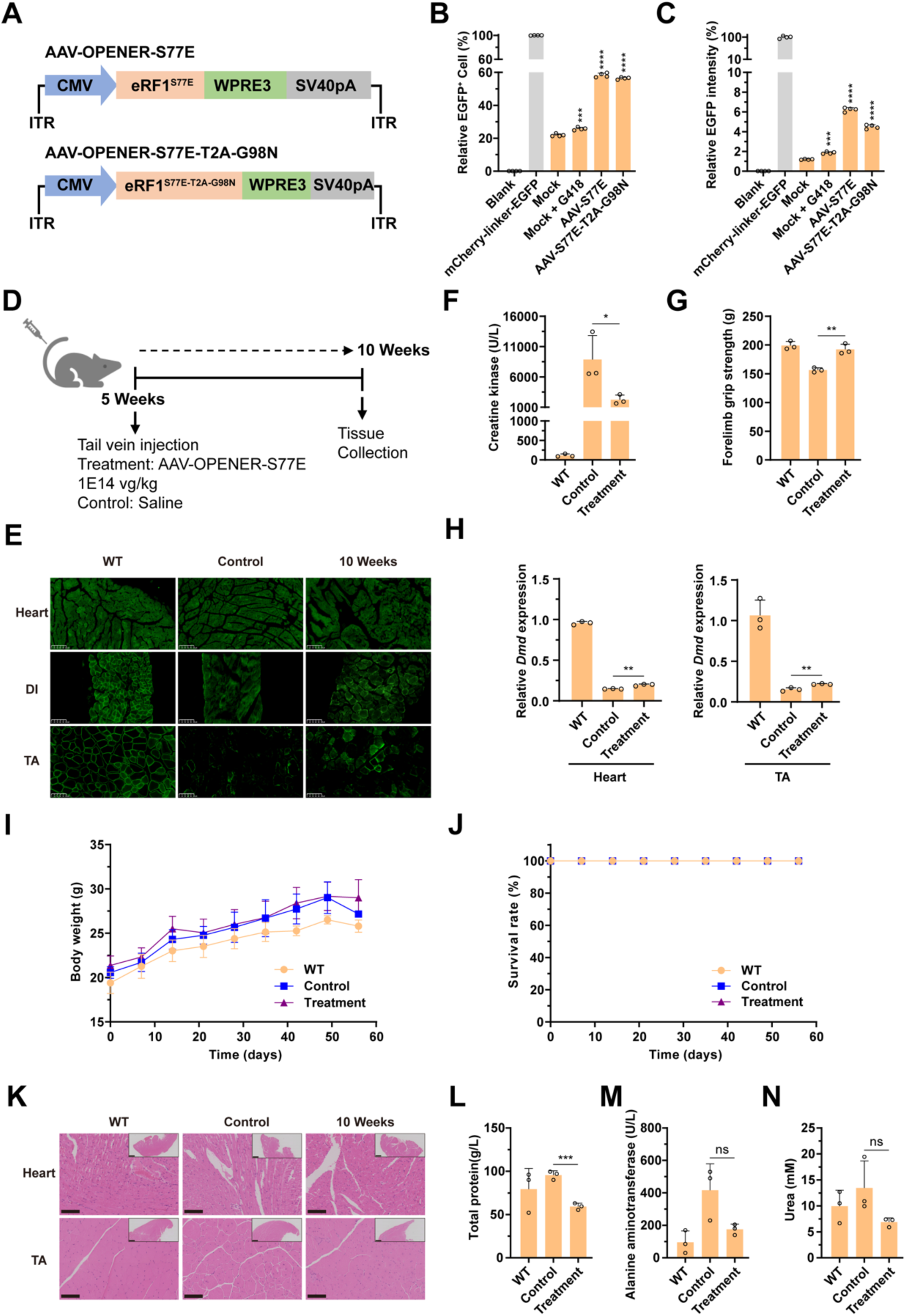
AAV-OPENER mediated restoration of dystrophin in vivo. (A) Schematic of the AAV vectors encoding OPENER-S77E and OPENER-S77E-T2A-G98N. (B-C) Flow analysis performed 72h after co-transfection of AAV-OPENER and Reporter-2 construct containing 60 nucleotides surrounding sequence of *Dmd* ^Q995TAA^ mutation (sequence in Supplementary fig.6A) into HEK293T cells, respectively. Bar plot showing the relative fraction of EGFP positive cells (B) and relative geometric mean fluorescence intensity of EGFP (C), normalized to that of the mCherry-linker-EGFP. Data are represented as the mean ± SD, n = 4. Mock and G418 treatments as controls. (D) Schematic of intravenous administration of AAV-OPENER-S77E particles. Tissues were collected for RNA analysis, immunofluorescence and H&E staining experiments at 10 weeks after treatment. Black arrows indicate time points for tissue collection after IV injection. (E) Immunofluorescence for dystrophin expression in tibialis anterior muscles (TA), diaphragm (DI), and heart of *Dmd*^Q995TAA^ mice was performed 10-week after IV injection. Dystrophin is shown in green. Scale bars, 100 μm. (F) Serum concentrations of creatine kinase were measured as a diagnostic marker of the muscle disease. (G) Forelimb grip strength was measured in Wild-type (WT) mice, *Dmd*^Q995TAA^ mice treated with PBS, and *Dmd*^Q995TAA^ treated with OPENER-S77E particles. (H) Quantification of *Dmd* mRNA by qPCR in heart and TA of *Dmd*^Q995TAA^ mice 10 weeks after injection. (I-J) Body weight curves (I) and survival rates (J) of indicated mouse groups within 56 days. Data are mean ± SD, n = 3 mice for all groups. (K) H&E staining of heart and TA after intravenous AAV treatment at 10 weeks. Scale bars, 100 μm; The top right insets show full images. Scale bars, 500 μm. (L-N), Serum biochemical analysis of total protein (L), alanine aminotransferase (M) and urea (N) in AAV-treated mice at 10 weeks, compared with wild-type mice, and PBS-treated *Dmd*^Q995TAA^ mice. Data are mean ± SD, n = 3 mice for all groups.

### Evaluation of in vivo safety of the AAV-OPENER

We next assessed the safety of the systemic AAV-OPENER delivery. No significant body weight loss or mouse death was observed within 56 days (Figs. 4I and 4J). H&E staining showed that partial correction of the muscle pathology of *Dmd*^Q995TAA^ mice treated with AAV-OPENER at 10 weeks and revealed no additional obvious histochemical changes (Fig. 4K). Serum biochemical analysis also showed modest improvements in the levels of total protein, alanine aminotransferase and urea in AAV-treated *Dmd*^Q995TAA^ mice at 10 weeks compared with wild-type C57BL/6 mice and *Dmd*^Q995TAA^ mice (Figs. 4L-N). Together, these data suggest that the in vivo safety of the AAV-OPENER system for potential therapeutic applications in DMD.

### OPENER suppress multiple UAG codons by multi-site incorporation of nsAAs

To investigate whether OPENER can suppress multiple nonsense mutants within single gene by combine with engineered tRNA synthase-tRNA pairs, we constructed a reporter sfGFP(TAG)_3_ in which three codons (positions 101, 133, and 150) were replaced with UAG (Fig. 5A)(*32*). In this reporter, full-length GFP fluorescence can only be restored when all three amber codons are simultaneously suppressed. We co-transfected OPENER with PylS/PylT pair(*47*) genetically encoded *N^ε^*-(*tert*-Butoxycarbonyl)-L-lysine (BocK) and sfGFP(TAG)_3_ into HEK293T cells, using eRF1^E55D^ as control. In the presence of BocK, flow cytometry analysis showed that efficiency of OPENER (S77E, G92T, G98N, T102D, K106V, I112D, T137C, S77E-IRES-G98N and S77E-T2A-G98N) in suppressing multiple UAG codons were 27.77%, 14.50%, 23.63%, 15.20%, 15.50%, 13.33%, 14.60%, 30.30%, 35.37%, respectively, which were substantially higher than that observed in the absence of OPENER (6.61%) (Fig. 5B and fig. S7A). Among them, the efficiencies of OPENER (S77E, G98N, S77E-IRES-G98N and S77E-T2A-G98N) were higher than that of eRF1^E55D^ (19.00%). Consistently, in the presence of BocK, sfGFP fluorescence was observed (Fig. 5C), and full-length sfGFP protein was detected by western blotting, indicating that OPENER-S77E showed the highest efficiency (Fig. 5D).

**Fig. 5.**
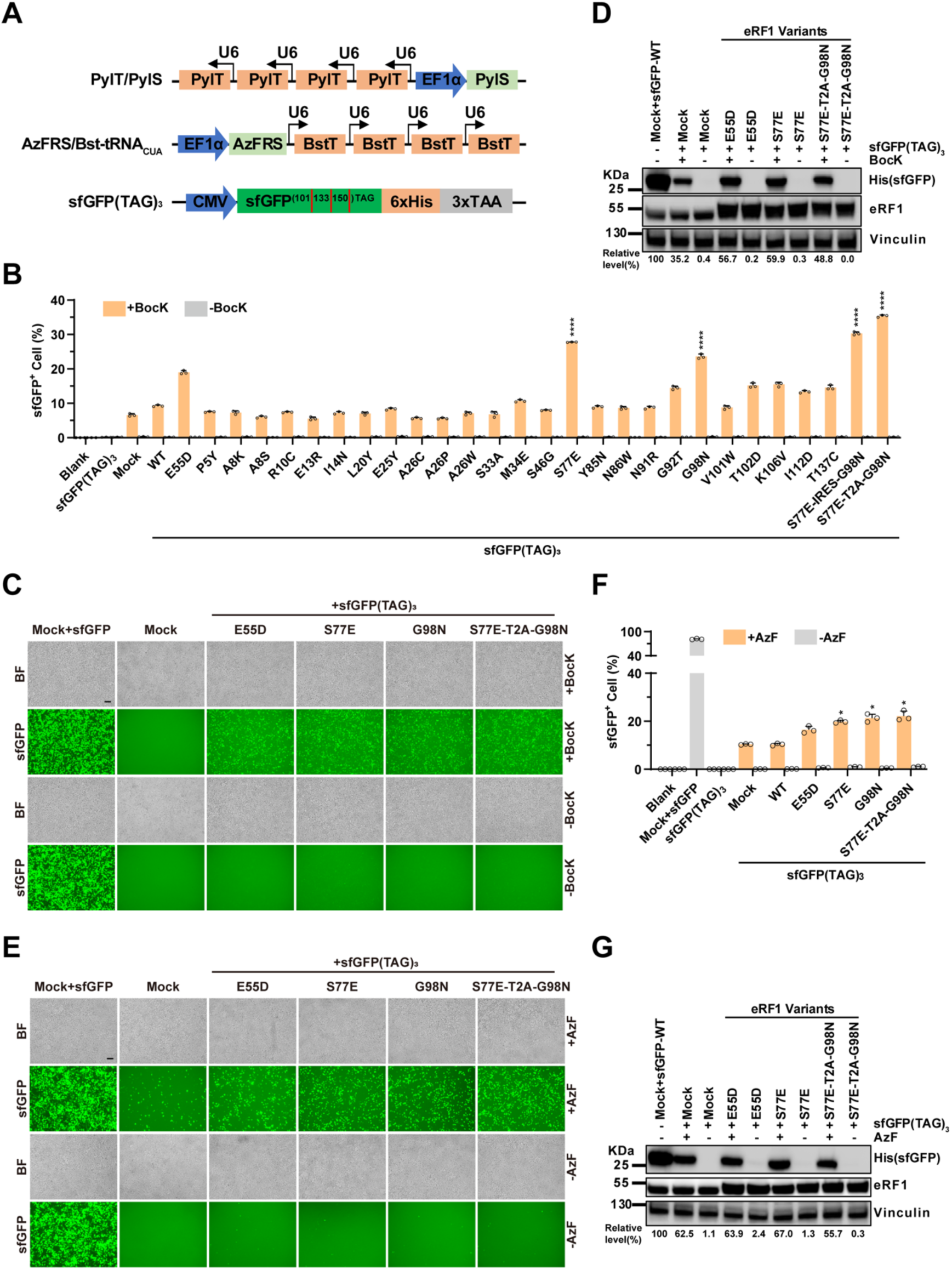
Multiple UAG nonsense codons were suppressed by OPENER with engineered tRNA synthase-tRNA pairs. (A) Schematic of the PylT/PylS system, the AzFRS/Bst-tRNA_CUA_ and sfGFP(TAG)_3_. (B-D) The PylT/PylS, sfGFP(TAG)_3_ and OPENER were transiently transfected into HEK293T cells and cultured in the presence or absence of 1 mM BocK for 72h. WT, wild-type eRF1 and eRF1^E55D^ served as controls. The percentage of sfGFP positive cells was quantified by flow cytometry (B). Data represent mean ± SD of three independent measurements. \**P* < 0.05; \*\**P* < 0.01; \*\*\**P* < 0.001; \*\*\*\**P* < 0.0001, compared to eRF1^E55D^ (two-sided Student′s *t*-test). Representative fluorescence images of cells showing the expression levels of sfGFP in the presence or absence of BocK (C), Scale bars, 100 μm. Western blotting analysis of the expression level of full-length sfGFP from cell lysates (D), the integrated optical density (IOD) of sfGFP bands was normalized to that of sfGFP-WT after load correction. Normalized IOD values are shown underneath the blot. (E-G) HEK293T cells were transiently transfected with the AzFRS/Bst-tRNA_CUA_, sfGFP(TAG)_3_ and OPENER, and grown in the presence or absence of 1 mM AzF for 72h. WT, Wild-type eRF1 and eRF1^E55D^ served as controls. Representative fluorescence images of cells showing the expression levels of sfGFP (E). Scale bars, 100 μm. The percentage of sfGFP positive cells was quantified by flow cytometry (F). Data represent the mean ± SD of three independent measurements. \**P* < 0.05; \*\**P* < 0.01; \*\*\**P* < 0.001; \*\*\*\**P* < 0.0001, compared to eRF1^E55D^ (two-sided Student′s *t*-test). The expression level of full-length sfGFP by western blotting from cell lysates (G), the integrated optical density (IOD) of sfGFP bands was normalized to that of sfGFP-WT after load correction. Normalized IOD values are shown underneath the blot.

We next tested the compatibility of OPENER with an alternative synthetase/tRNA pair. Co-transfection of OPENER (S77E, G98N and S77E-T2A-G98N) with sfGFP(TAG)_3_ and AzFRS/Bst-tRNA_CUA_ pair(*36*) genetically encoded *p-azido*-L-phenylalanine (AzF) into HEK293T cells (Figure 5A). In the presence of AzF, sfGFP fluorescence was observed (Fig. 5E) and flow cytometry analysis showed that efficiency of OPENER (S77E, G98N and S77E-T2A-G98N) were 19.83%, 21.33%, 22.17% respectively, all of which were higher than that of eRF1^E55D^ (16.47%) (Fig. 5F and fig. S7B). The full-length sfGFP protein was also observed by western blotting in the presence of AzF, indicating that OPENER-S77E showed the highest efficiency (Fig. 5G). Together, these results demonstrate that OPENER in combination with engineered tRNA synthase-tRNA pairs enables highly efficient suppression of multiple TAG codons in the presence of nonstandard amino acids, achieving multisite-specific incorporation of nonstandard amino acids into single protein, with OPENER-S77E showing the highest efficiency.

### Enhanced BocK incorporation in conditional eRF1-knockout cells

The OPENER may compete with endogenous eRF1 to limit nsAAs incorporation(*32*). To investigate whether endogenous eRF1 affect OPENER mediated nsAAs incorporation, we generated conditional eRF1-knockout HEK293T cells by site-specific integration of the Tet-off *ETF1* coding-sequence cassette into the AAVS1 safe-harbor locus via CRISPR/Cas9-mediated homology-directed repair and subsequently disrupting the endogenous *ETF1* alleles through CRISPR/Cas9 targeting of the exon–intron junction to avoid cleavage within *ETF1* coding sequence (Fig. 6A and figs. S8A, S8B). In this cell line, the endogenous eRF1 is completely depleted after 7d of doxycycline treatment, whereas wild-type eRF1 is overexpressed when cultured in medium without doxycycline, as validated by western blotting (Fig. 6B). We evaluated BocK incorporation at multiple UAG sites by co-transfecting OPENER with PylS/PylT pair and sfGFP(TAG)_3_ into wild-type HEK293T cells and conditional eRF1-knockout cells separately. In the presence of BocK, sfGFP fluorescence was detected (Fig. 6C) and flow cytometry analysis showed that the efficiencies of OPENER-S77E, OPENER-G98N, eRF1^E55D^ and control (without OPENER) were 29.07%, 26.23%, 25.80%, and 8.58% in wild-type HEK293T cells (Fig. 6D), and the corresponding efficiencies were higher, reaching 32.80%, 30.93%, 27.03%, and 13.70% in conditional eRF1-knockout cells cultured in medium with doxycycline (Fig. 6E). By contrast, in conditional eRF1-knockout cells cultured in medium without doxycycline, in which wild-type eRF1 remained overexpressed, corresponding efficiencies decreased to 6.85%, 4.28%, 6.06% and 1.21% (Fig. 6F). Notably, OPENER-S77E exhibited the highest efficiency and the highest geometric mean fluorescence intensity under eRF1-knockout condition (Fig. 6D-F).

**Fig. 6.**
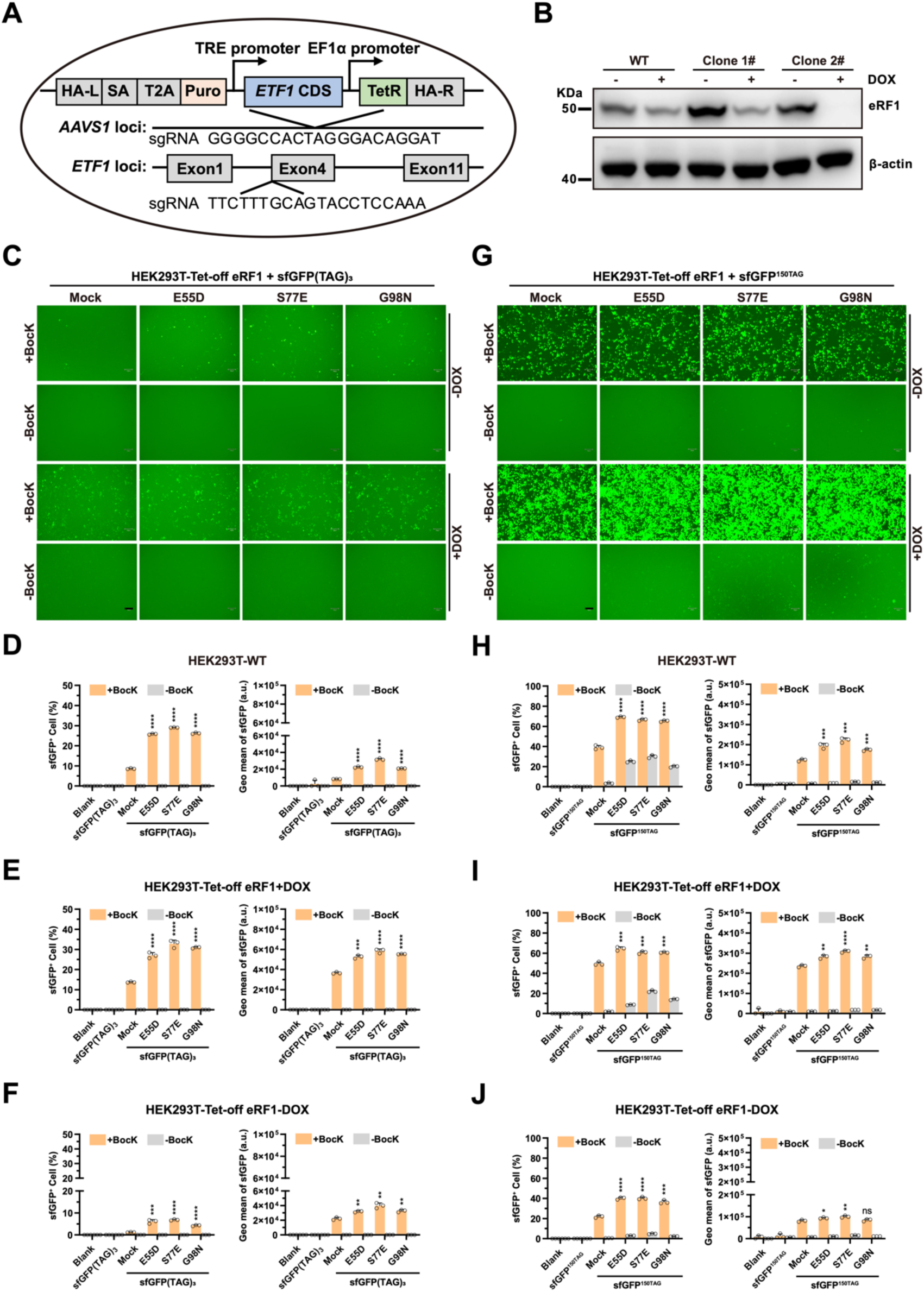
Evaluation of incorporation efficiency of BocK in conditional eRF1-knockout cells. (A) Schematic of Dox-inducible eRF1-knockout HEK293T cells. Using CRISPR/Cas9-mediated homology-directed repair, we first achieved site-specific integration of the Tet-off ETF1 coding-sequence cassette into the AAVS1 safe-harbor locus and then disrupted the endogenous ETF1 alleles by CRISPR/Cas9 targeting of the exon-intron junction, avoiding cleavage within exogenous ETF1 coding sequence. (B) Western blotting analysis of expression of eRF1 protein in each clone of Dox-inducible eRF1 knockout HEK293T cells 7 days after adding doxycycline in cell culture medium. (C) Representative fluorescence images of cells showing the expression levels of sfGFP after co-transfection of sfGFP(TAG)_3,_ PylT/PylS and OPENER in Dox-inducible eRF1 knockout cells cultured with doxycycline or without doxycycline, and grown in the presence or absence of 1 mM Bock for 72h, separately. Scale bars, 100 μm. (D-F) sfGFP(TAG)_3,_ PylT/PylS and OPENER were transiently transfected into in wild-type HEK293T cells(D), and Dox-inducible eRF1 knockout cells cultured with doxycycline (E) or without doxycycline (F), and grown in the presence or absence of 1 mM BocK for 72h, separately. The percentage of sfGFP positive cells and geometric mean fluorescence intensity of sfGFP was quantified by flow cytometry. Data represent the mean ± SD of three independent measurements. \**P* < 0.05; \*\**P* < 0.01; \*\*\**P* < 0.001; \*\*\*\**P* < 0.0001, compared to Mock (two-sided Student′s *t*-test). (G) Representative fluorescence images of cells showing the expression levels of sfGFP after co-transfection of sfGFP^150TAG^, PylT/PylS and OPENER in Dox-inducible eRF1 knockout cells cultured with doxycycline or without doxycycline, and grown in the presence or absence of 1 mM BocK for 72h, separately. Scale bars, 100 μm. (H-J) sfGFP(TAG), PylT/PylS and OPENER were transiently transfected into in wild-type HEK293T cells(H), and Dox-inducible eRF1 knockout cells cultured with doxycycline (I) or without doxycycline (J), and grown in the presence or absence of 1 mM BocK for 72h, separately. The percentage of sfGFP positive cells and geometric mean fluorescence intensity of sfGFP was quantified by flow cytometry. Data represent the mean ± SD of three independent measurements. \**P* < 0.05; \*\**P* < 0.01; \*\*\**P* < 0.001; \*\*\*\**P* < 0.0001, compared to Mock (two-sided Student′s *t*-test).

We next co-transfected OPENER with PylS/PylT pair and sfGFP^150TAG^ into wild-type HEK293T cells or conditional eRF1-knockout cells separately. In the presence of BocK, sfGFP fluorescence was detected (Fig. 6G) and flow cytometry analysis showed that efficiencies of OPENER-S77E, OPENER-G98N, eRF1^E55D^ and control (without OPENER) were 66.50%, 65.33%, 69.26% and 39.07% in wild-type HEK293T cells (Fig. 6H), the corresponding efficiencies were 60.70%, 60.73%, 64.47% and 49.53% in conditional eRF1-knockout cells cultured in medium with doxycycline (Fig. 6I), whereas the efficiencies decreased to 40.13%, 36.43%, 40.10% and 21.93% in conditional eRF1-knockout cells cultured in medium without doxycycline (Fig. 6J). Notably, OPENER-S77E exhibited the highest geometric mean fluorescence intensity, although percentage of sfGFP positive cells were slightly lower than that of eRF1^E55D^ (Fig. 6H-J). Collectively, these results demonstrate that OPENER combined with engineered PylS/PylT pair systems efficiently suppresses one and multiple UAG codons in wild-type cells and further enhances BocK incorporation in conditional eRF1-knockout cells cultured in the presence of doxycycline. Conversely, the markedly reduced efficiencies observed in conditional eRF1-knockout cells cultured in the absence of doxycycline suggest that stable overexpression of wild-type eRF1 may compete with OPENER, thereby limiting nsAAs incorporation.

## Discussions

Here, we demonstrate that OPENER is a powerful tool for efficient suppression of nonsense mutations. Both OPENER-S77E alone and combined with OPENER-G98N can efficiently suppress all three types of stop codon in multiple cell lines and also suppress 12 different nonsense mutants across four different genes in reporters, including *IDUA, CFTR, DMD and RPE65*, which lead to four different genetic diseases. These findings provide a generalizable strategy for efficiently suppressing nonsense mutations across multiple disease contexts, with the potential to extend to many other diseases caused by nonsense mutations. Existing approaches, including small-molecule readthrough compounds and some engineered suppressor tRNAs usually show limited efficacy depending on strong sequence context of PTCs(*48–51*). OPENER offer an approach to suppress PTCs through targeted reprogramming translation termination via engineered eRF1, avoiding broad ribosome perturbation and cytotoxicity commonly associated with small-molecule readthrough compounds (*52–54*). Hence, OPENER system provides a promising disease-agnostic therapeutic approach for effective and safe treatment of rare genetic diseases caused by nonsense mutations.

Although the precise structural mechanism by which the S77E and G98N substitutions promote readthrough requires further elucidation, we can speculate based on the known architecture of human eRF1. Serine 77 is located in a region connecting the NIKS motif and the GTS loop, both critical for stop codon decoding. Substitution with a negatively charged glutamate (S77E) may alter the local electrostatic landscape or conformational dynamics of the N-domain, potentially relaxing its stringency for stop codon recognition. Glycine 98, due to its minimal side chain, often confers flexibility. Mutation to asparagine (G98N) might introduce a stabilizing hydrogen bond or restrict flexibility, thereby indirectly affecting codon interaction. The synergistic effect observed in the combined variant suggests these mutations may act through complementary, non-identical pathways to compromise termination fidelity.

Our results show that systemic AAV-mediated delivery of OPENER-S77E in *Dmd*^Q995TAA^ mice restored dystrophin expression and ameliorated muscle pathology, providing compelling in vivo proof of concept for therapeutic application. DMD arises from deletions, duplications or point mutations in the *DMD* gene, which encodes dystrophin, a large intracellular protein in muscle cells that maintains membrane integrity during contraction. Notably, ∼10% by point mutations and more than 50% are nonsense mutations that introduce PTCs into the *DMD* gene (*55, 56*). Full-length of *DMD* cDNA (∼14kb) far extends the package capacity of AAV, and gene replacement is constrained by vector size, potential immunogenicity, and transgene-related toxicities. OPENER overcome these limitations by restoring endogenous dystrophin translation without requiring delivery of the full-length coding sequence. An aminoacyl-tRNA-synthase (aaRS)/tRNA pair was reported to readthrough a nonsense mutation in *mdx* mice(*57*), but delivery, readthrough efficiency and bioavailability of nsAAs is limited. Besides, UAA mutations represent particularly challenging substrates for existing therapeutic strategies. Readthrough compounds exhibit poor efficiency on UAA codons(*50*), adenine base editors struggle to correct such sites due to low efficiency and bystander editing (*58*), and UAA is an unfavorable substrate for ADAR-mediated RNA editing (*9, 59*). AAV-mediated delivery of OPENER therefore represents a valuable addition to therapeutic genome-editing.

OPENER also offers a versatile chemical tool in synthetic biology. When combined with engineered aminoacyl-tRNA synthetase/tRNA pairs, OPENER-S77E enabled efficient suppression of multiple UAG codons, and achieved site-specific incorporation of non-standard amino acids BocK and AzF into single protein separately in human cells in the presence of nsAAs. The efficiency of OPENER-S77E was higher than previously described eRF1 mutant E55D in response to the three amber stop codons(*32, 37*). Moreover, OPENER-S77E with engineered tRNA synthase-tRNA pairs systems showed the highest suppression efficiency of multiple UAG codons in conditional eRF1-knockout cells cultured in medium with doxycycline, further enhancing nsAAs incorporation. Conversely, in the absence of doxycycline, the markedly reduced efficiency observed for OPENER-S77E suggests that overexpression of wild-type eRF1 may compete with OPENER. Mechanistically, OPENER may relieve competition with endogenous eRF1 to combine with eRF3(*20, 60*), thereby promoting readthrough of UAG codon by engineered aminoacyl-tRNA synthetase/tRNA pairs. Thus, OPENER expands the synthetic biology toolkit for precisely controlling translation and engineering therapeutic proteins.

Our study has several important limitations. First, the efficiency of OPENER mediated suppression of nonsense should be evaluated across more proteins in cells and disease models in vivo. Second, off-target effects were not observed in vivo for long time. Third, the mechanism of OPENER mediates suppression of nonsense mutants need to be fully elucidated, although OPENER may be potentially competitive interactions with endogenous eRF1. Finally, the clinical efficacy of readthrough-promoting OPENER will depend on additional parameters such as targeted delivery and tissue specificity.

Taken together, our findings demonstrate that OPENER efficiently suppresses nonsense mutations in cells, and that AAV delivery of OPENER can rescue gene function and ameliorate disease phenotypes in vivo, providing proof-of-concept for treating DMD. OPENER also suppresses multiple UAG codons within a single gene by enabling efficient multisite-specific incorporation of nsAAs in human cells when combined with engineered aminoacyl-tRNA synthetase/tRNA pairs, further enhancing nsAAs incorporation in conditional eRF1-knockout cells cultured in the presence of doxycycline. The OPENER system offers a promising universal strategy for treating multiple nonsense mutation diseases and expands the synthetic biology toolbox for precise control of protein translation and efficient production of modified therapeutic proteins.

## Acknowledgements

We thank Longlong Si and Xiaoshan Shi for helpful discussions and Shenzhen Synthetic Biology Infrastructure support with facilities. This work was partially supported by grants from National Key R&D Program of China (No.2022YFC3400200 to Y.C.) and National Natural Foundation of China (No.32101173 to Y.C.). C.L. was supported by National Natural Science Foundation of China (Nos. 32488301, 32025022, 32230062) and Strategic Priority Research Program of Chinese Academy of Sciences (No. XDB0480000). M.L. was supported by National Natural Science Foundation of China (No.32300798)

## Author Contributions

Y.C., C.L. and G.C. conceived and supervised the study. Y.C. designed experiments with input from C.L. and G.C.; Y.C., Z.Z., Y.Q., and M.L. performed experiments and analyzed the data. Y.X. conducted most of the bioinformatics analysis. P.G. prepared for some plasmids and cell culture. Y.C. wrote the manuscript with input from C.L., G.C., and all other authors.

