## Supplementary Material for "Engineered translation release factor 1 suppresses disease-causing nonsense mutations and enables multi-site nonstandard amino acids incorporation"

### Materials and Methods

**eRF1 variants library and plasmids construction.** The Homo sapiens eRF1 coding gene (Transcript ID: ENST00000360541.10) and saturation library targeting N-terminal domain of human eRF1(amino acids 2-142) were synthesized by GenScript. For the saturation library, each amino acid within residues 2-142 was individually replaced with the other 19 amino acids. All wild-type and variant eRF1 fragments were cloned into plasmid (Addgene 52962). The sequence encoding wild-type sfGFP, sfGFP$^{150TAG}$, sfGFP$^{150TGA}$, sfGFP$^{150TAA}$ and sfGFP$^{101,\ 133,\ 150TAG}$ were synthesized by GenScript as previously reported(*1*). The positive reporter mCherry-linker-EGFP was synthesized by GenScript and cloned into pGenleti plasmid (from GenScript)， and the mCherry-linker-EGFP vector was digested with BamHI and NheI to generate the backbone for mCherry-PTC-EGFP reporters, followed by gel purification using Gel extraction kit (Zymo Research cat# 11-301C). Annealed oligos (Table S9) as fragments were ligated into the digested mCherry- linker -EGFP vector using T4 DNA ligase (NEB cat# M0202S) at 16 °C overnight, separately. After the ligation reaction, 2 µl each reaction was transformed into a competent *E.coli* strain TransStbl3 chemically competent cell (Transgen cat# CD521-01) according to the manufacturer's instructions. Plasmid DNA was isolated using a Plasmid Plus Midi Kit (25) (QIAGEN cat# 12943) according to the manufacturer's instructions.

All PCR amplifications were performed using Q5 High-Fidelity 2X Master Mix (NEB cat# M0494S) or 2×TransStart® FastPfu PCR SuperMix (TransGen cat# AS231-01). The PylS/ PylT pair plasmids was obtained from Jason chin lab (*2*) and AzFRS/Bst-tRNA$_{CUA}$ pair plasmids was generated as described previously by ourselves (*3*). All enzymes and buffers were obtained from New England Biolabs unless otherwise noted. Nuclease-free water (Life Technologies cat# 10977-015) was used for all cloning and PCR reactions. All gene fragments in this paper were synthesized by GenScript unless otherwise noted. All primers and oligos were synthesized and sanger sequencing by Sangon Biotech.

**Amplification of Lentivirus library.** We performed the amplification of Lentivirus plasmid library, following the guidelines (https://www.addgene.org/protocols/pooled-library-amplification/) with minor modifications, Briefly, 200 µL *E.coli* HST08 Premium Electro-Cells (TakaRa cat# 9028) was divided into eight aliquots of 25 µL for electroporation. For each aliquot, 100 ng of plasmid DNA was added and mix gently before electroporation. Electroporation was performed using MicroPulser TM (Bio-Rad cat#1652100) in 0.2 cm cuvette, with Ec2, 2.5 kV, 1 pulse. Immediately, 1 mL SOC was added to cuvette and transferred into 14 mL vented falcon tube containing 3 mL SOC after pulsing treatment. This process was repeated for all eight aliquots. A new tube for two electroporation and the four transformation tubes were incubated at 30-37 °C with shaking at 225 rpm for 1 hour. After recovery, all cultures were pooled and mixed thoroughly. The sequential 1:100 dilutions of the cells (add 10 μL of the pool to 990 μL LB then perform a second and then third 1:100 dilution), and 100 μL of each dilution was plated on pre-warmed LB agar plates. Plates were incubated at 30 °C overnight. Next morning, colonies were counted on the most dilute Petri dish. Total colony yield = 3000 × 100 × 100 × 100 / 0.1, The total number of recovered colonies was confirmed to be at least 1,000-fold greater than the number of perturbations in the library. For harvesting, agar plates were kept on ice. Cold LB (100 mL) was prechilled at 4 °C, and four or more sterile 50 mL conical tubes were placed on ice. Bacterial lawns were scraped using a sterile spreader while repeatedly washing plates with 10 mL ice cold LB per scrape, pooling two plates per 50 mL conical tube. Scraping was repeated until the agar surface was clear. Approximately 25 mL of LB bacterial suspension was obtained per plate. Cells were pelleted by centrifugation at 4,000 × g, 4 °C, 15 min, and supernatant was removed. Pellets were weighed, and plasmid DNA was purified using the Qiagen HiSpeed Maxi Kit (Qiagen cat# 12662) following the manufacturer's protocol. DNA concentrations were quantified using a Nanodrop (Thermo Fisher).

**Cell culture.** HEK293T, Neuro-2a and NIH3T3 cell lines were from ATCC. HEK293T cells, Neuro-2a cells and NIH3T3 cells were maintained in high-glucose Dulbecco's Modified Eagle Medium (Corning cat# 10-013-CV) with 10% (v/v) fetal bovine serum (FBS, Gibco cat# 10082147) and 1% Penicillin-Streptomycin (Hyclone cat# SV30010), cells were cultured at 37 °C with 5% $CO_2$ and passaged every 3-4 days, and tested for mycoplasma with Universal Mycoplasma Detection Kit (ATCC® 30-1012K™) every 4-6 weeks.

**Lentiviral library packaging.** Lentiviral particles were produced in HEK293T cells using a standard three plasmid packaging system. HEK293T cells were seeded into 10 cm dishes and maintained in high-glucose DMEM. When cultures reached 70-80% confluence, the medium was replaced with 10 mL fresh complete DMEM. For each dish, a transfection mixture was prepared by combining 10 μg pMD2.G, 10 μg psPAX2, 10 μg Lenti-library plasmid, and 60 μL TransIT-X2® transfection reagent (Mirus, cat# MIR 6000) in Opti-MEM. After gentle pipette mixing, the mixture was incubated at room temperature for 15-30 min before being added dropwise to the cells. Cultures were gently rocked and returned to the incubator. At 24 h post-transfection, 5 mL fresh medium was added. At 48 h, supernatants were harvested into 50 mL tubes, and 10 mL medium was replenished. Supernatants were centrifuged at 3,000 rpm for 15 min, and then filtered through 0.45 μm filters, and stored at 4 °C. At 72 h, a second supernatant harvest was collected, and cells were briefly rinsed with PBS to recover residual virus. The combined supernatants were filtered as above. The viral supernatants were transferred into sterile 39 mL Beckman Quick-Seal tubes. Remaining volume was filled with cold medium, leaving a 2-3 mm gap below the seal. Tubes were sealed using a heat sealer and balanced to ≤ 0.02 g. Ultracentrifugation was performed at 25,000 rpm for 2.5 h at 4 °C using a pre-cooled rotor. Following centrifugation, pellets were identified visually and the supernatant was carefully discarded. Viral pellets were gently resuspended in 500 μL cold PBS (Corning cat# 21-040-CVC), and each tube was rinsed with another 500 μL PBS to maximize recovery. Care was taken to avoid bubble formation to prevent viral loss. All resuspended material was pooled, aliquoted in 20 μL volumes into pre-chilled 0.2 mL tubes, and stored at -80 °C.

**Lentiviral titer determination.** Lentiviral titers were measured following the procedures described in reference(*4*), with minor modifications. Frozen viral aliquots were thawed on ice. HEK293T cells were digested, counted, and $2 \times 10^6$ cells were transferred into 1.5 mL tubes. The volume was adjusted to 1 mL, and 8 μg/mL polybrene (Beyotime, cat# C0351) was added. A virus dilution series (0, 1, 2, 4, 8, 16, and 32 μL virus) was added to the cells, mixed gently, and transferred to 6-well plates, bringing the final volume to 2 mL per well. After 24 h, cells were digested and counted. For viability-based titer determination, $4 \times 10^3$ cells per condition were seeded into black, clear-bottom 96-well plates. For each virus dilution, 8 replicate wells were prepared, 4 wells received 100 μL standard medium, 4 wells received 100 μL medium containing 10 μg/mL Blasticidin HCl (Beyotime, cat# ST018). Additionally, 4 control wells received medium only. Plates were incubated for 3 days. Viability measurements were performed using the CellTiter-Glo® Luminescent Cell Viability Assay (Promega cat# G7571). The CellTiter-Glo buffer and lyophilized substrate were equilibrated to room temperature, reconstituted according to the manufacturer's protocol, and 100 μL reagent was added to each well. Plates were shaken for 2 min, incubated for 10 min, and luminescence was recorded. Infectious units per milliliter were calculated as: $IU/mL = 10^3 \times N \times P / V$, where: N = number of cells at the time of transduction; P = fraction of antibiotic-resistant (positive) cells; V = volume (μL) of virus added. The fraction P was calculated by subtracting background viability from the untreated control, followed by normalization to viability in non-selective wells.

**Lentivirus infection and single cell clonal isolation.** HEK293T cells were transduced with the lentiviral library at a low multiplicity of infection (MOI ~ 0.1) in the presence of 8 μg/mL polybrene. At 72 h post-infection, selection was initiated by supplementing the medium with 10 μg/mL blasticidin and maintained for 7 days to enrich for stably integrated cells. For single-cell clonal isolation, blasticidin resistant cells expressing green fluorescent protein (GFP) were single-cell sorted using fluorescence-activated cell sorting (FACS) into flat bottom 96-well plates containing 100 μL DMEM supplemented with 10% FBS and 1% Penicillin-Streptomycin per well. Plates were incubated for 10-14 days, with medium changes as needed, until wells with well-defined, expanding single-cell colonies were identified. For expansion, individual colonies were detached by adding 20 μL Accutase (STEMCELL Technologies, cat# 07920) per well and neutralizing with 20 μL growth medium. Cells were then transferred into 24-well plates containing 800 μL medium for further propagation.

**Genomic DNA extraction.** For NGS, DNA was extracted using the EasyPure Genomic DNA Kit (TransGen cat# EM101-02) according to the manufacturer's protocol. For Sanger sequence, cells were washed with PBS, and lysed in containing 200 μL of QuickExtract™ DNA Extraction Solution (Lucigen Cat# QE09050) per well of 48-well plates, and genomic DNA (gDNA) was extracted using the manufacturer's protocol. Briefly, the sorted plates were sealed, vortexed and heated at 65°C for 6 min then 98°C for 2 min.

**NGS sequence and frequency analysis of eRF1 variants.** Genomic DNA extracted from each single-cell clone was used as the template for PCR amplification of the eRF1 variant region. Target fragments were amplified using Q5 High-Fidelity 2× Master Mix (NEB cat# M0494S) according to the manufacturer's instructions. PCR products were assessed by agarose gel electrophoresis, and the correct-sized bands were excised and purified using the EasyPure® Quick Gel Extraction Kit (Omaga cat# D2500-02). Purified amplicons were submitted to GENEWIZ (Azenta Life Sciences) for next-generation sequencing (NGS) library preparation and sequencing. Primer sequences used for PCR amplification and library construction are listed in Table S9. The raw fastq data were processed using fastp with default parameters to remove adapter sequences and low-quality reads. The processed reads were then aligned to the coding DNA sequence (CDS) of ETF1 gene using Bowtie 2(*5*) under default parameters for paired-end alignment. Samblaster (*6*) with parameters --excludeDups and --addMateTags was employed to filter out PCR duplicates and abnormally aligned reads, yielding reads effectively mapping to the ETF1 CDS region. Due to paired-end sequencing, the filtered valid reads were assembled into fragments based on their alignment coordinates and overlapping sequences. Subsequently, all fragments within the target region were subjected to multiple sequence alignment against the CDS of ETF1 as reference sequence using BLASTN(*7*). The optimal alignment results were used to determine the base identity at each position within the target region for every fragment. The insertion and deletion events in the alignment results were processed according to the optimal alignment strategy for subsequent sequence alignment, and counts of these events are excluded. The nucleotide sequences were then translated into codon lists by grouping every three bases. The frequency and proportion of each codon type at each position were calculated. All significant mutation codons with a frequency exceeding 2% were identified as potential ETF1 mutation types present in the sample. Finally, the number of occurrences and the average frequency of the identified codon mutations across all samples were summarized for subsequent analysis and experimentation.

**Transient transfection, imaging and flow analysis.** Transfections were performed using either the TransIT-X2® Dynamic Delivery System (Mirus, cat# MIR 6000) or Lipofectamine 3000 (Thermo Fisher Scientific, cat# L3000001) following the manufacturers' protocols with minor modifications. One day prior to transfection, $1.4 \times 10^5$ cells were seeded per well in 24-well plates containing 500 μL complete medium and allowed to adhere overnight. For co-transfection of individual OPENERs plasmids and reporter plasmids, a total of 200 ng DNA per well was used, consisting of 100 ng base editor plasmid and 100 ng sfGFP Reporter-1s or Reporter-2s plasmid. Transfection complexes were prepared in Opti-MEM (Gibco cat#31985070) using 0.6 μL TransIT-X2 or Lipofectamine 3000 per well, incubated 20 min at room temperature, and added dropwise to cells. At the indicated time points, fluorescent protein–positive cells were assessed by fluorescence microscopy (Thermo EVOS). Cells were then dissociated, pelleted by centrifugation at 800 rpm for 3 min, resuspended in complete medium, and subjected to flow cytometry (Beckman CytoFLEX S) to quantify sfGFP or EGFP reporter protein expression by CytExpert software or FlowJo.

**Western blotting.** HEK293T cells were transfected with either 6×His-tagged Reporter-1 constructs (TAA, TAG, or TGA) or 6×His-tagged Reporter-2 constructs in combination with

OPENER-S77E or OPENER-S77E-T2A-G98N, as indicated. Cells were harvested 72 h post-transfection and lysed in RIPA buffer (Beyotime cat# P0013C) supplemented with protease and phosphatase inhibitor cocktails (MedChemExpress cat# HY-K0010). Total protein concentrations were determined using the BCA Protein Assay Kit (Beyotime, cat# P0012). For each sample, 20 μg of total protein was resolved on 4-12% FuturePAGETM gels 12-well (ACE cat# ET12412Gel) and transferred onto PVDF membranes at 160V for 50 min. Membranes were blocked in 10% (w/v) nonfat milk (Beyotime, cat# P0216) in TBST (Beyotime, cat# ST673) for 2 h at room temperature. Blots were incubated overnight at 4 °C with the following primary antibodies (all at 1:1,000 dilution): anti-actin (ABclonal, cat# AC026), anti-His (Bioss cat# bsm-330044M), anti-eRF1 (Abcam cat# ab31799), anti-Vinculin (MedChemExpress cat# HY-P80372). The next day, membranes were washed using TBST and then incubated with HRP-conjugated Goat Anti-mouse IgG (H+L) secondary antibody (Beyotime, cat# A0216) or HRP-conjugated Goat Anti-Rabbit IgG (H+L) secondary antibody (Beyotime, cat# A0208) for 1 h at room temperature. Protein bands were visualized using enhanced chemiluminescence (Beyotime, cat# P0018AS) and imaged with a UVP ChemStudio 815 imaging system (Analytikjena).

**Mass spectrometry for immunoprecipitation and peptide identification.** HEK293T cells were transfected with His-tagged Reporter-1 constructs (TAG, TGA, or TAA) together with OPENER-S77E or OPENER-S77E-T2A-G98N) separately. 72 h After transfection, sfGFP positive cells were isolated by fluorescence-activated cell sorting (FACS) and lysed on ice for 20 min in lysis buffer containing 50 mM $NaH_2PO_4$, 300 mM NaCl, pH 7.4, and protease inhibitor cocktail (MedChemExpress cat# HY-K0010). Lysates were clarified by ultracentrifugation at 12000 rpm for 15 min at 4 °C. The resulting supernatant was incubated with Ni-charged MagBeads (GenScript cat# L00295) at 4 °C for 5 h. Beads were washed four times with 1mL wash buffer (50 mM $NaH_2PO_4$, 300 mM NaCl, 10 mM imidazole, pH 7.4), followed by elution using 500 μL Elution Buffer (50 mM $NaH_2PO4$, 300 mM NaCl, 250 mM imidazole, pH 7.4) for 2 h at 4 °C. Eluted protein complexes were resolved by SDS-PAGE and visualized by Coomassie Brilliant Blue staining. Gel bands corresponding to full-length Reporter-1 (TAA, TAG, TGA) were excised and destained in 25 mM ammonium bicarbonate / 50% acetonitrile. Following dithiothreitol (DTT) reduction and iodoacetamide (IAA) alkylation, proteins were digested overnight at 37 °C with sequencing-grade trypsin (Pierce). Tryptic peptides were extracted with acetonitrile containing 0.2% formic acid (FA), dried in a vacuum concentrator at

30 °C, and resuspended in 15 μL 0.1% FA, quantified by Nanodrop. LC–MS/MS analysis was performed on a Bruker timsTOF Pro 2. Spectra were searched using DIANN against the Human UniProt database and a custom database that included the sequences with 20 different "X" amino acids in the position corresponding to the 150th site in the CDS region of sfGFP. Only the peptides across or downstream of the 150th amino acid of sfGFP were used to calculate the abundance of sfGFP protein. Peptide abundances were quantified based on precursor ion intensities, and the relative frequency of amino acid insertion at the 150th residue of sfGFP was calculated from the normalized abundance of corresponding peptides.

**Quantitative proteome analysis by mass spectrometry.** HEK293T cells were co-transfected with His-tagged Reporter-1 (TAG, TGA, or TAA) and OPENER-S77E and OPENER-S77E-T2A-G98N) constructs separately. 72 h after transfection, sfGFP positive cells were sorted by FACS and were digested with RIPA lysis buffer (Beyotime cat# P0013C). The digested peptide was dissolved in 0.1% formic acid (FA)/$H_2O$. Then, the peptide was quantified by Nanodrop, and the concentration was adjusted to 200 ng/μL. 2 μg peptide was used for mass spectrometry analysis (a Bruker timsTOF Pro 2). The MS raw files were processed with timsControl software, and spectra were searched by DIANN against the Human UniProt database.

**Animal use.** All experiments involving live animals were approved by the Institutional Animal Care and Use Committee and the Animal Ethics Committee of Cyagen Biosciences. All mouse experiments were performed at Cyagen Biosciences Inc (Project Number: OMN250717HS1). For experiments involving *Dmd*$^{Q995TAA}$ mice strain from Cyagen Biosciences. Wild-type mouse and *Dmd*$^{Q995TAA}$ mice were C57BL/6J strains. All mice were kept in a specific-pathogen-free animal room with temperatures controlled at 20-26 °C and relative humidity controlled at 40–70% under a 12 h dark-light cycle access to standard rodent diet and water.

**AAV construction and delivery.** AAV-OPENER-S77E and AAV-OPENER-S77E-T2A-G98N were constructed and packaged in AAV8 by PackGene Biotech. The titer of AAV-PylRS/tRNA vector was 1 × $10^{14}$ GC/ mL. Tail-vein injections in mice were performed under sterile conditions. AAV were diluted to a final dose of 1 × $10^{14}$ vg/kg prior to administration. Mice were restrained in a dedicated mouse restrainer, and the tail was gently extended to ensure full immobilization. To enhance vein visibility, the tail was warmed for 1-2 min using a heating lamp or by brief immersion in warm water. The central lateral tail vein was identified under illumination, and the injection site was disinfected with an alcohol swab. A 0.5 mL syringe fitted with a 27-gauge needle was inserted at a shallow 10-20° angle, with the needle oriented parallel to the vein. The vector solution was injected slowly, and proper intravenous placement was confirmed by the absence of resistance or swelling; injections were halted immediately if extravasation was suspected. Following administration, the needle was withdrawn smoothly and a sterile cotton swab was applied to the injection site to prevent bleeding. Mice were monitored post-injection to ensure normal recovery and to detect any adverse reactions.

**Histological analysis.** The tibialis anterior muscle and heart tissues of different mice were isolated and fixed in 4% PFA solution for two days, sequentially dehydrated with 75%, 85%, 90%, 95% and 100% ethanol, and defatted with xylene for 2 h before being embedded in paraffin. The 5-μm-thick sections were cut and subjected to H&E staining (Hematoxylin, Baso cat# BA4097; Eosin, MERCK cat# E4382). The slices were observed under optical microscopy, and the histological morphology of different mouse groups was compared.

**Immunofluorescence staining.** Tibialis anterior (TA), heart, and diaphragm tissues from different mice, which were fixed in 4% PFA at room temperature for 1 h, and sequentially cryoprotected in 10%, 20%, and 30% (w/v) sucrose solutions at 4 °C, transferring to the next concentration after the tissue had sunk. Cryoprotected tissues were blotted dry, oriented in embedding molds with fresh OCT (Sakura), frozen at -20 °C for 30 min, and stored at -80 °C until sectioning. Frozen OCT blocks were equilibrated in the cryostat (Leica) for 30 min, and sections of the desired thickness were collected and stored at -20 °C until use. For staining, sections were permeabilized in 0.5% Triton X-100(Sigma cat# 93443) in PBS (ThermoFisher cat# 10010023) for 15 min, washed three times in PBS, and blocked in 5% BSA (Sigma cat# A9418) for 1 h at room temperature. Sections were incubated with anti-dystrophin (1:500, Abcam cat# ab15277,) diluted in blocking buffer overnight at 4 °C, followed by a 30 min equilibration at room temperature and three PBS washes (3 × 5 min). Fluorescent secondary antibodies and DAPI (Invitrogen cat# 62248) were applied for 1 h at room temperature in the dark, after which slides were washed three times in PBS and mounted using an antifade mounting medium for imaging under a Nikon Ti-S microscope.

**Grip strength test.** Grip strength was assessed at day 80 to evaluate forelimb muscle strength using a grip-strength meter (Shanghai Xinruan XR501). Prior to testing, the grip-strength meter was thoroughly cleaned to remove residual odors and debris, disinfected with alcohol, and calibrated according to the manufacturer's instructions. Each animal was identified and recorded before measurement. Mice were gently guided toward the horizontal grasping bar, allowing them to grip the bar voluntarily without applying excessive force. A consistent backward pull was applied until the mouse released its grip, and the instrument automatically recorded the peak tension. Each animal underwent three consecutive trials, and the mean value was used for analysis.

**qPCR analysis.** Total RNA from heart or mouse tibialis anterior (TA) muscle from different mice was isolated using using TRIGene reagent (Genstar cat# P118) following a modified phenol-chloroform extraction procedure. RNA concentration and purity were measured using a NanoDrop 2000 spectrophotometer, and aliquots were stored at -80 °C to avoid repeated freeze–thaw cycles. Purified RNA was reverse-transcribed into cDNA and subjected to quantitative real-time PCR (qPCR) using 2×RealStar Fast SYBR qPCR Mix (Genstar cat# A301-05) and cycling conditions recommended. Gene expression levels were normalized to GAPDH as an internal control, and relative transcript abundance was calculated using the $2^{-\Delta\Delta Ct}$ method. Primer sequences used in this study are listed in Table S9.

**Serum biochemical analysis.** Peripheral blood was collected from mice at 10 weeks via retro-orbital bleeding under sterile conditions. Prior to the procedure, sterile lancets, serum collection tubes, 70% ethanol, and hemostatic materials were prepared. Mice were lightly anesthetized with isoflurane to minimize pain and stress, then gently restrained to maintain a stable posture. A sterile capillary lancet was inserted into the retro-orbital venous sinus, allowing blood to flow directly into collection tubes. Blood samples were allowed to clot at room temperature or at 4 °C for 30 min, followed by centrifugation at 1,500-2,000 × g for 10-15 min to separate serum. The clarified supernatant was carefully transferred into sterile tubes and stored immediately at -20 °C or -80 °C until analysis. Serum biochemical parameters, including creatine kinase (CK), alanine aminotransferase (ALT), blood urea nitrogen (BUN), and total protein (TP) were measured using standard clinical chemistry assays. Three mice were used in each group.

**Generation of conditional eRF1 knockout cells.** To generate stable cell lines, $1 \times 10^6$ HEK293T cells were seeded into each well of a 6-well plate 24 h prior to transfection. Cells were first co-transfected with 3 μg PX458-sgRNA targeting the AAVS1 locus and 1 μg donor plasmid containing the Tet-off cassette (synthesized by GenScript) using Lipofectamine 3000 according to the manufacturer's instructions. At 72 h post-transfection, cells were subjected to puromycin selection (2 μg/mL) for 7–10 days to obtain polyclonal pools stably harboring the Tet-off cassette. Next, the puromycin-resistant HEK293T pool was transfected with 3 μg PX458-sgRNA targeting the endogenous eRF1 locus. After 3-5 days, GFP-positive cells were isolated via flow cytometry and single cells were sorted into 96-well plates for clonal expansion. Individual clones were then expanded in 24-well plates and screened by genomic DNA extraction, PCR amplification, gel electrophoresis, and next-generation sequencing (NGS) to confirm correct insertion and editing at the target locus. Long-term cultures were periodically tested to ensure continued puromycin resistance.

**BocK and AzF incorporation.** HEK293T cells were seeded 18-24 h prior to transfection at a density of $1.4 \times 10^5$ cells per well in 24-well plates. For Tet-off eRF1 stable cell lines, cells were maintained in medium with or without doxycycline (1 μg/mL) for 5 days prior to seeding. Cells were then plated 18–24 h before transfection at a density of $1.4 \times 10^5$ cells per well in 500 μL of medium containing doxycycline (1 μg/mL) or doxycycline-free medium. Cells were cultured until they reached ≥70% confluence, with 0.5 mL complete medium added per well. For nonstandard amino acid (nsAA) supplementation, BocK (TCI cat# B1670) or AzF (MedChemExpress cat# HY-16714) was added directly to the designated wells immediately before transfection to a final concentration of 1 mM. Transfection complexes (50 μL Opti-MEM, 100 ng of each plasmid, and 0.6 μL TransIT-X2 per well) were prepared in RNase-free 1.5-mL tubes, followed by gently mixing and a 20-min incubation at room temperature. The transfection mixture was added gently onto the surface of the culture medium. Plates were gently rocked 6–8 times, and returned to the incubator for further culture. At 72 h post-transfection, cells were imaged under a fluorescence microscope to observe sfGFP positive cells (nsAAs incorporation efficiency), and representative images were acquired. Cells were then dissociated, centrifuged at 800 rpm for 3 min, resuspended in complete medium, and subjected to flow cytometry analysis (Beckman CytoFLEX).

**Statistics and reproducibility.** All statistical analyses were performed on at least three biologically independent experiments using GraphPad prism10. Error bars represent the mean ± SD. Detailed information on exact sample sizes and experimental replicates can be found in the individual figure legends. Tests for statistically significant differences between groups were performed using a two-tailed Student's *t*-test and $*P < 0.05$; $**P < 0.01$; $***P < 0.001$; $****P < 0.0001$ were considered significant.

**Data availability.** Sequencing data have been deposited in the NCBI Sequence Read Archive database with accession code GSE313505. All plasmids in this study will be available upon reasonable request.

**Code availability.** Codes have been uploaded to the Github repository https://github.com/0CBH0/amplicon_handle, including code and files for reproducibility.

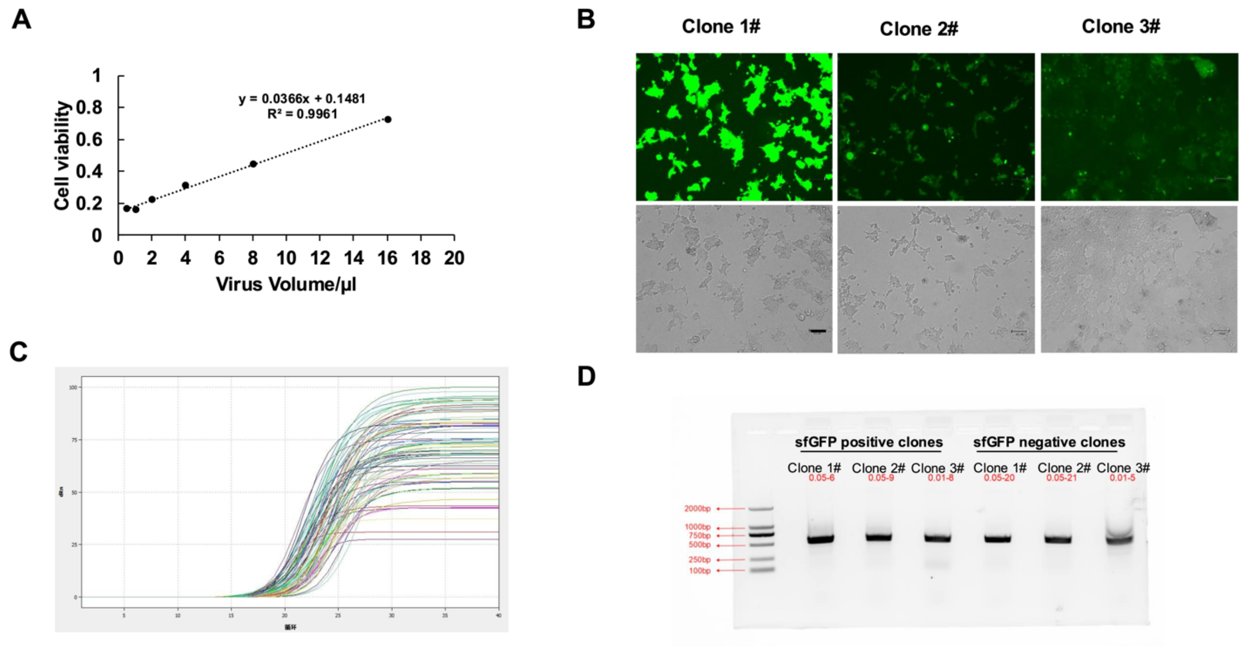


**Supplementary Figure 1. A saturated mutagenesis library screen targeting the N-terminal domain of human eRF1.** (A) The titer curve of lentivirus library. Cell viability and virus volume when detection of virus titer. (B) Representative fluorescence images of single cells showing the expression levels of sfGFP during library screening. Scale bars, 100 µm. (C) Determination of PCR cycles number by qPCR during of NGS PCR. 30 cycles can avoid PCR overamplification. (D) Representative gel image of NGS PCR for CDS sequence of the N-terminal domain of human eRF1.

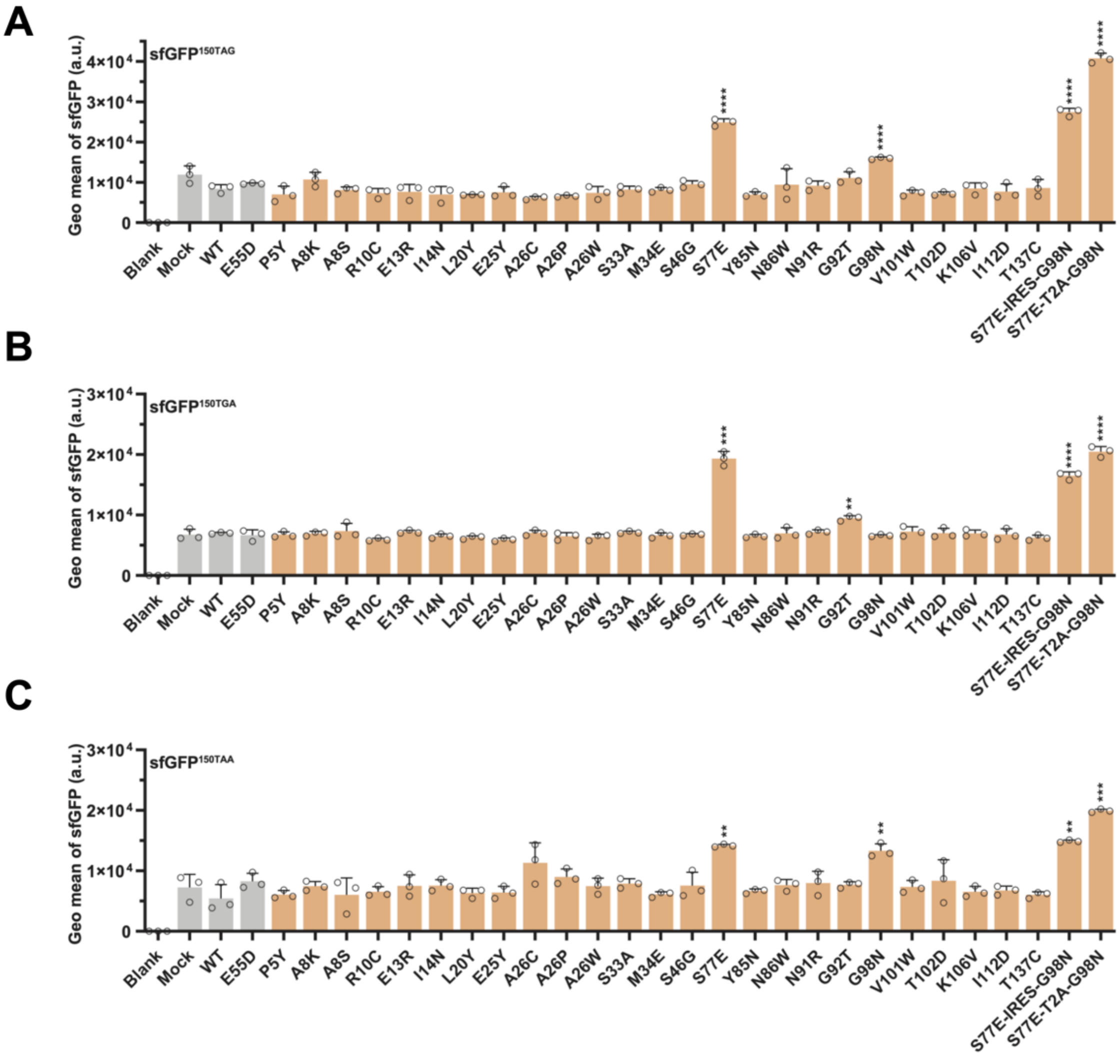


**Supplementary Figure 2. Validation of OPENER-mediated suppression of nonsense mutations in HEK293T cells.** (A-C) Flow analysis was performed 72h after co-transfection of Reporter-1s construct, sfGFP$^{150TAG}$ (A), sfGFP$^{150TGA}$ (B), sfGFP$^{150TAA}$ (C), with OPENER separately. Bar plot shows the geometric mean intensity of sfGFP positive cells. WT, wild-type eRF1 and eRF1$^{E55D}$ served as controls. Data are mean ± SD, n = 3; **P* < 0.05; ***P* < 0.01; ****P* < 0.001; *****P* < 0.0001, compared to eRF1$^{E55D}$ (two-sided Student′s *t*-test).

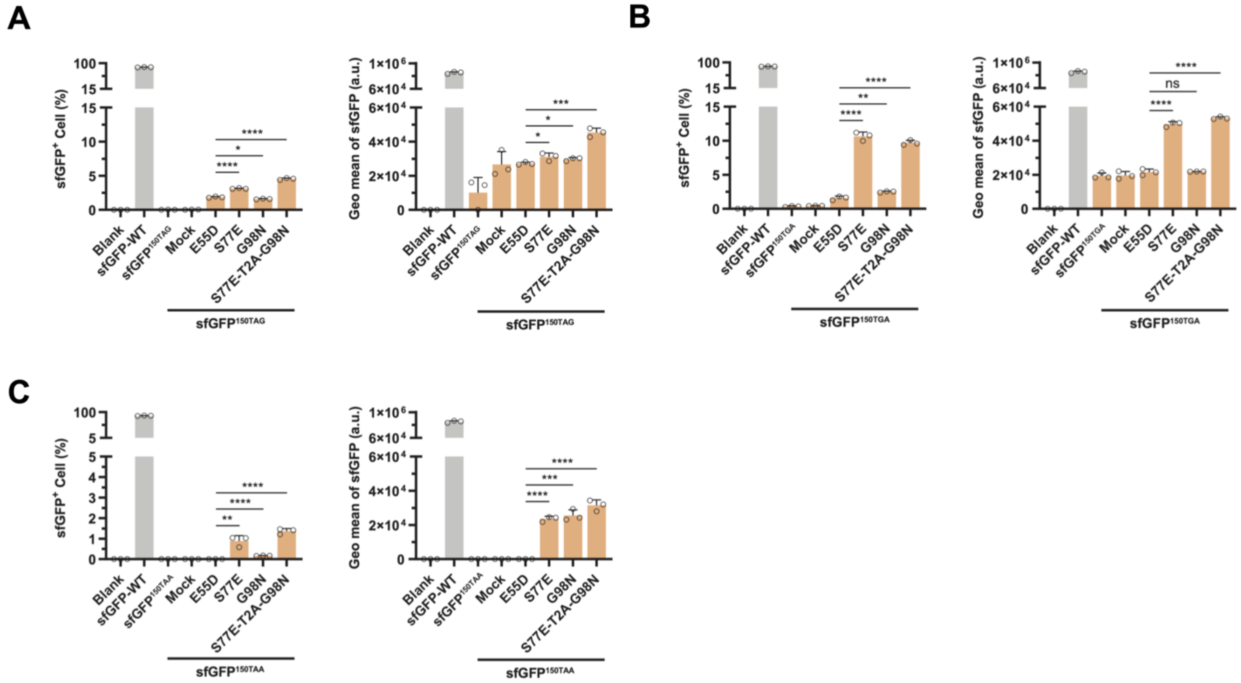


**Supplementary Figure 3. Detection of OPENER-mediated suppression of nonsense mutations in Neuro-2a cells. (A-C)** Flow analysis of the percentage of sfGFP positive cells and Geo MFI intensity of sfGFP when 72h after co-transfection of OPENER with Reporter-1s, $sfGFP^{150TAG}$ (A), $sfGFP^{150TGA}$ (B), $sfGFP^{150TAA}$ (C), into Neuro-2a cells separately. Data are mean ± SD, n = 3; *$P < 0.05$; **$P < 0.01$; ***$P < 0.001$; ****$P < 0.0001$, compared to $eRF1^{E55D}$ (two-sided Student′s *t*-test).

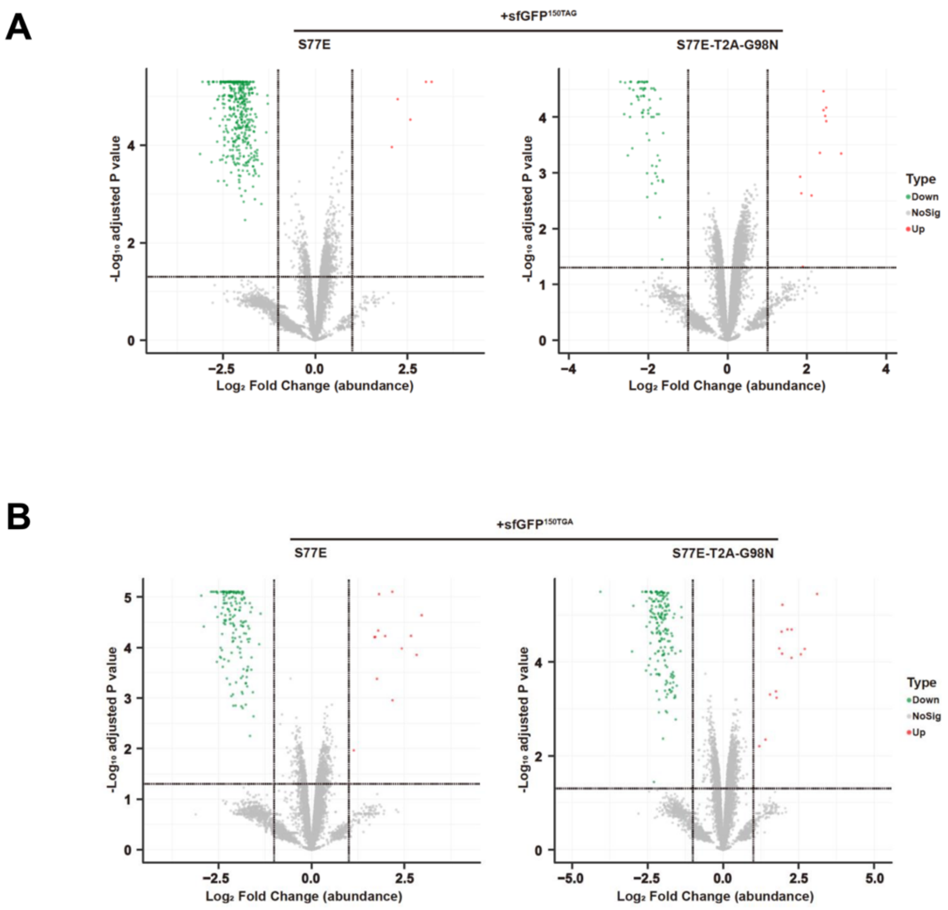


**Supplementary Figure 4. Evaluation of potential effect of OPENER on the proteome.** (A-B) Protein-level analysis of whole proteome mass spectrometry data from sfGFP positive cells sorted by FACS after co-transfection of OPENER-S77E and OPENER-S77E-T2A-G98N with $sfGFP^{150TAG}$ and $sfGFP^{150TGA}$ reporter separately. $Log_2$ fold change in protein abundance compared to sfGFP-WT control cells. Dashed lines indicate an adjusted *p*-value of 0.05 on the y-axis and $log_2$ fold change in abundance of ±1 compared with control cells.

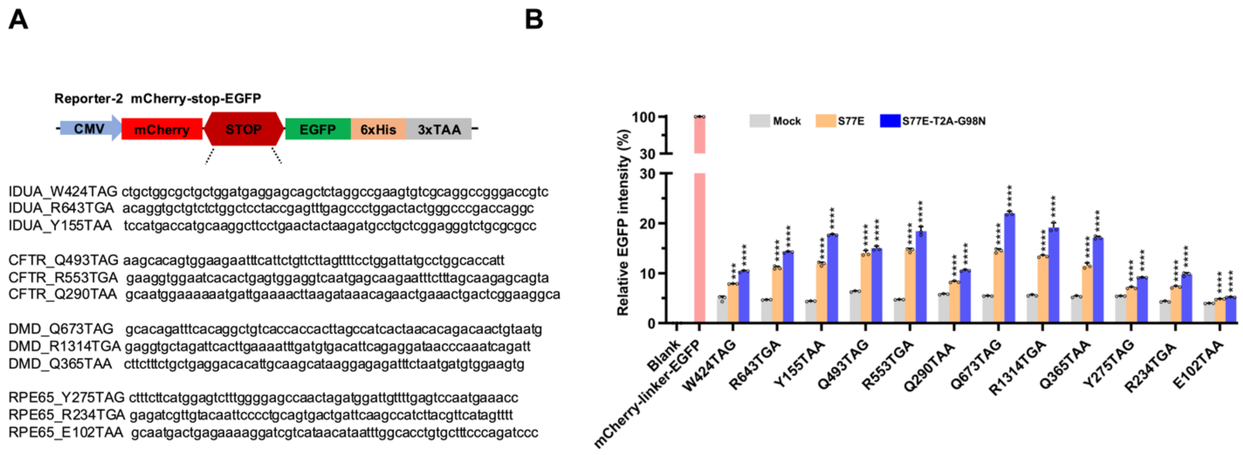


**Supplementary Figure 5. OPENER suppress the pathogenic PTCs.** (A) Schematic of the Reporter-2s constructs. Each pathogenic PTC with 60 nucleotides (nts) of surrounding sequence context was inserted into a dual fluorescent protein reporter. Full sequences of 60 nucleotides are shown. (B)geometric mean intensity of sfGFP was quantified by flow cytometry 72h after co-transfection of OPENER-S77E and OPENER-S77E-T2A-G98N and Reporter-2s constructs were transfected into HEK293T, respectively. Bar plot shows the geometric mean intensity of EGFP positive cells normalized to that of the mCherry-linker-EGFP. Data are represented as the mean ± SD, n = 3; **P* < 0.05; ***P* < 0.01; ****P* < 0.001; *****P* < 0.0001, compared to Mock (two-sided Student′s *t*-test).

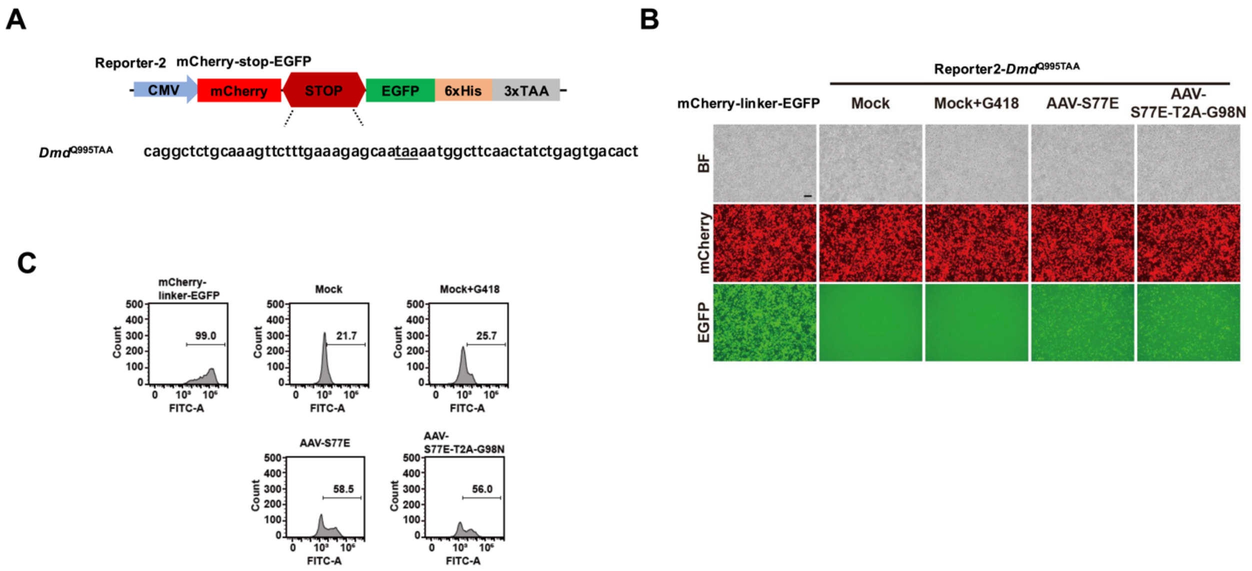


**Supplementary Figure 6. Quantifying readthrough of mouse *Dmd*$^{Q995TAA}$ in reporter.** (A) Schematic of the Reporter2-*Dmd*$^{Q995TAA}$ constructs. (B) Representative fluorescence images of cells showing the expression levels of mCherry-EGFP 72h after AAV-OPENER-S77E and AAV-OPENER-S77E-T2A-G98N with Reporter2-*Dmd*$^{Q995TAA}$ constructs were transfected into HEK293T, respectively. Scale bars, 100 µm. Mock and G418 treatments as controls. (C) Flow cytometry confirmed the expression of mCherry-EGFP in HEK293T cells 72h after AAV-OPENER-S77E and AAV-OPENER-S77E-T2A-G98N with Reporter2-*Dmd*$^{Q995TAA}$ constructs were transfected into HEK293T, respectively. The y axes denote the cell count, and the x axes denote the EGFP fluorescence intensity. The positive rate of cells is shown. Mock and G418 treatments as controls.

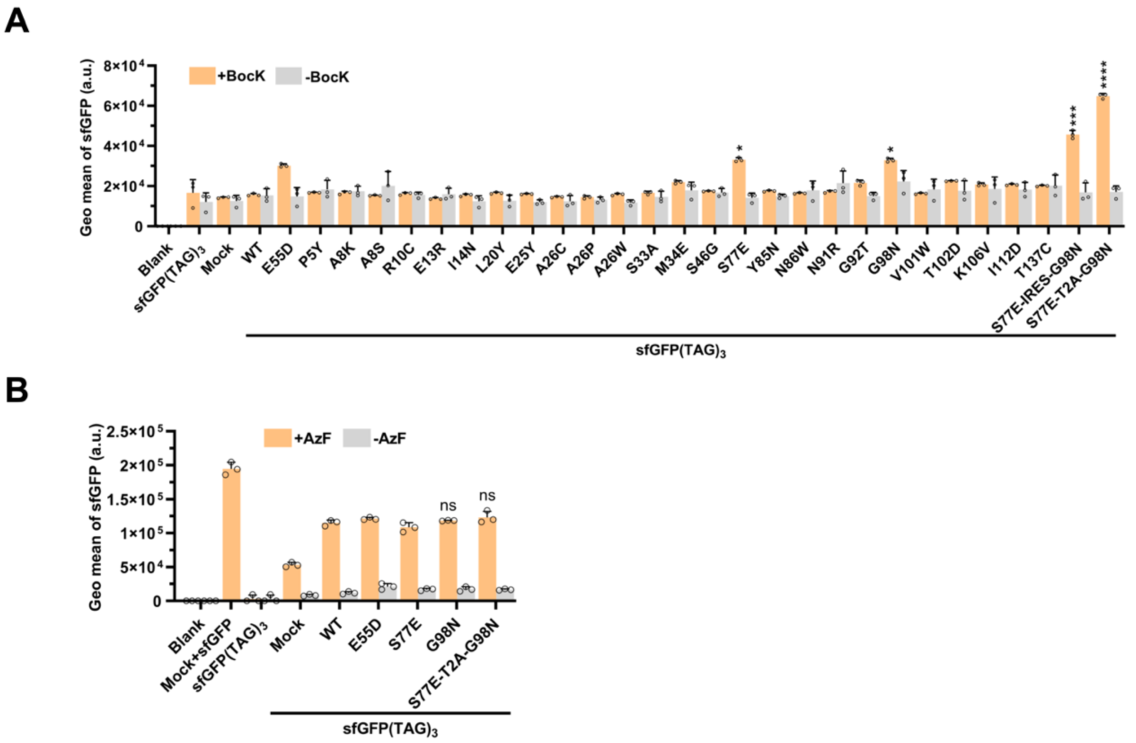


**Supplementary Figure 7. OPENER with engineered tRNA synthase-tRNA pairs enable suppress multiple UAGs nonsense.** (A) The PylT/PylS, sfGFP(TAG)$_3$ and OPENER were transiently transfected into HEK293T cells, and cultured in the presence or absence of 1 mM BocK for 72h. WT, wild-type eRF1 and eRF1$^{E55D}$ served as controls. The geometric mean intensity of sfGFP positive cells was quantified by flow cytometry. Data represent mean ± SD of three independent measurements. *$P < 0.05$; **$P < 0.01$; ***$P < 0.001$; ****$P < 0.0001$, compared to eRF1$^{E55D}$ (two-sided Student′s *t*-test). (B) HEK293T cells were transiently transfected with the AzFRS/Bst-tRNA$_{CUA}$, sfGFP(TAG)$_3$ and OPENER, and grown in the presence or absence of 1 mM AzF for 72h. The geometric mean intensity of sfGFP positive cells was quantified by flow cytometry.

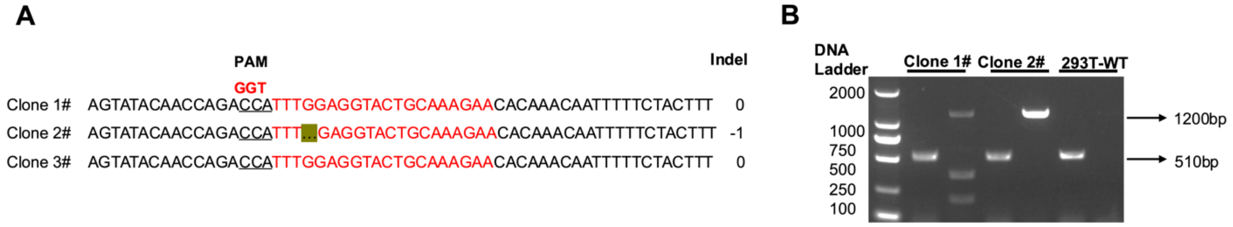


**Supplementary Figure 8. Generation of conditional eRF1-knockout HEK293T cell line.** (A)NGS analysis after genome extraction of single clone showing the sfGFP positive cells were sorted by flow cytometry after 3-5 days of transfection of 3ug PX458-sgRNA targeting endogenous eRF1, grown in 24 wells. (B) Gel running after PCR amplify for knockin tet-off cassette from single clones. Top band (1200 bp) for knock-in Tet-off cassette and down band (510 bp) for wild-type allege band in Clone 2#.

#### Supplementary tables S1 to S9

Table S1. Frequency and proportion of eRF1 variants in different clones.

Table S2. Evaluation of amino acids incorporated at the 150th residue of sfGFP by OPENER suppress three different stop codons.

Table S3. Quantitative proteomics analysis of OPENER-S77E on sfGFP(TAA) _DiffExp.

Table S4. Quantitative proteomics analysis of OPENER-S77E on sfGFP(TAG) _DiffExp.

Table S5. Quantitative proteomics analysis of OPENER-S77E on sfGFP(TGA)_DiffExp.

Table S6. Quantitative proteomics analysis of OPENER-S77E-T2A-G98N on sfGFP(TAA)_DiffExp.

Table S7. Quantitative proteomics analysis of OPENER-S77E-T2A-G98N on sfGFP(TAG)_DiffExp.

Table S8. Quantitative proteomics analysis of OPENER-S77E-T2A-G98N on sfGFP(TGA)_DiffExp.

Table S9. Oligos and sanger sequence primers used in this study.
